# Quality control of HLA-DR molecules by the lysosomal aspartyl protease, cathepsin D

**DOI:** 10.1101/2020.03.25.004101

**Authors:** Nakul Shah, Sarah McDonald, Charles T.A. Parker, Ryan W.M. Robinson, Amin Oomatia, Alessandra De Riva, Michael J. Deery, Elizabeth D. Mellins, Robert Busch

**Affiliations:** Dept of Life Sciences, University of Roehampton, London, UK; Dept of Medicine, University of Cambridge, Cambridge, UK; Cambridge Centre for Proteomics, Cambridge, UK; Dept of Pediatrics, Program in Immunology, Stanford University, Stanford, CA, USA

**Keywords:** Major histocompatibility complex class II, human leukocyte antigen DR, lysosomal proteolysis, aspartyl cathepsins, protein turnover, quality control, peptide binding, mass spectrometry, flow cytometry, Western blot.

## Abstract

Major histocompatibility complex class II (MHCII)^1^ molecules display peptides on antigen-presenting cells (APCs) for inspection by CD4+ T cells. MHCII surface levels and life span are regulated post-translationally by peptide loading and ubiquitin-dependent lysosomal targeting, but proteases responsible for MHCII protein degradation remain unidentified. Here, we examined the role of aspartyl proteases in MHCII protein degradation and characterised the form of MHCII that is degraded. Exposure of immature monocyte-derived dendritic cells (MoDCs) and KG-1 acute myeloid leukaemia cells to the aspartyl protease inhibitor, pepstatin A (PepA), caused accumulation of human leukocyte antigen (HLA)-DR molecules in intracellular vesicles. Statistically significant PepA effects on MHCII protein expression were also observed in murine APCs. In KG-1 cells, cathepsin D (CatD) was the sole expressed aspartyl protease, and shRNA-mediated knockdown ablated the PepA effect, providing formal proof of CatD involvement. *In vitro*, CatD initiated specific cleavage of recombinant DR at αF54, a site flanking the peptide-binding groove. Immunochemical characteristics of PepA-rescued DR molecules in KG-1 cells were consistent with selective CatD attack on HLA-DR molecules that lack association with the MHCII chaperone, invariant chain, or with stably bound peptide. We propose that CatD has a critical role in the selective lysosomal disposal of mature HLA-DR molecules that have lost, or never acquired, bound peptide, explaining how MHCII protein life span is coupled to peptide loading.

## Introduction

Major histocompatibility complex class II (MHCII) glycoproteins present diverse self and foreign peptides to drive immune tolerance and adaptive immunity, respectively, after detection by antigen receptors of CD4+ T lymphocytes (1). Under the control of the class II transactivator (CIITA), MHCII molecules are expressed constitutively on myeloid antigen-presenting cells (APCs), B lymphocytes and thymic epithelial cells and induced on other cell types by proinflammatory cytokines (2, 3). Peptide loading of MHC class II molecules is largely confined to endosomes, due to regulation of intracellular transport and occupancy of the peptide-binding groove by invariant chain, a chaperone that binds MHCII molecules in the endoplasmic reticulum (4–6). Nascent MHCII αβ dimers, after assembling with Ii, are transported to late endosomes/early lysosomes, where proteases successively trim Ii, leaving class II-associated Ii peptides (CLIP) in the peptide-binding groove. CLIP is then exchanged for endosomal peptides, either spontaneously or through catalysis by HLA-DM, a non-classical MHCII homolog that stabilises empty MHCII molecules, protects them from degradation, and edits in favour of kinetically stable peptide/MHCII complexes. These complexes are expressed at the plasma membrane and, after a time, the MHCII molecules may be re-internalised into lysosomes and degraded. One unresolved question is how MHCII molecules can survive in the proteolytic environment of endocytic compartments while being loaded with peptides, only to be degraded later when needed.

MHCII protein turnover rates are affected by the cellular milieu, with half-lives ranging from hours to weeks (7). *In vivo*, MHCII degradation rates are slower in murine B cells than dendritic cells (DCs) (8). In human and murine DCs differentiated *in vitro*, MHCII degradation is shut down by proinflammatory cytokines and pathogen-derived molecular patterns (9, 10): in non-activated DCs, ubiquitylation of the MHCII β-chain cytoplasmic tail (catalysed by the E3 ligase, MARCH-1) promotes internalization and enables lysosomal degradation, whereas MARCH-1 expression ceases when DCs are activated (11–14). This mechanism may allow the persistent presentation of antigen by DCs during CD4+ T-cell activation and exhaustion (15, 16). In follicular B cells and macrophages, downregulation of constitutively expressed MARCH-1 is involved in the induction of MHCII by interleukin-10 (17, 18), whereas at the centroblast stage of B-cell differentiation in the germinal center, induction of MARCH-1 and MHCII internalization limits antigen presentation during the selection of mutated BCRs for high antigen affinity (19).

Another factor regulating the turnover of MHCII proteins is their peptide loading state. Peptide dissociation half-lives limit the persistence of peptide/MHCII protein complexes on APC surfaces (20). Acquisition of a stable peptide repertoire can be impaired in cells that lack the peptide editor, DM, and/or due to un-edited peptide loading following spontaneous CLIP release from MHCII allelic variants with low CLIP affinity; in such settings, MHCII protein turnover is accelerated (21–25). Biophysical studies show that tightly-bound peptides stabilise MHCII molecules (26, 27), yet the molecular mechanisms linking poor peptide loading to lysosomal MHCII protein turnover *in vivo* remain undefined.

Little is known about the proteases that mediate physiological MHCII turnover. Early biochemical studies highlighted the resistance of tightly folded MHCII ectodomains to proteases *in vitro* (28, 29), but the enzymes used in these studies are not available during lysosomal protein turnover *in vivo*. Inhibition of lysosomal cysteine proteases rescued only a small proportion of the MHCII protein, human leukocyte antigen (HLA)-DR, on the surface of human immature monocyte-derived dendritic cells (MoDCs)(30); moreover, *in vitro* digestion of purified DR with several cysteine proteases failed to produce detectable cleavage (31). The serine protease, cathepsin (Cat) G, cleaved HLA-DR β-chains *in vitro*, but a specific CatG inhibitor had no effect *in vivo*, so the relevance to physiological degradation is doubtful (31). Genetic screens for factors influencing MHCII protein expression failed to identify any lysosomal proteases (32). Similarly, targeted knockouts of various cathepsins in mice provided no evidence of specific involvement in the turnover of MHCII αβ dimers, despite clear evidence that several cathepsins are involved in processing Ii (33) or in the generative or destructive processing of antigenic peptides (34, 35).

Here, we examined the roles of two aspartyl proteases found in APCs, cathepsin D and cathepsin E (CatD/E) in MHCII protein turnover. These enzymes use active-site aspartyl residues to mediate proteolytic attack on peptide bonds and can be blocked with a specific inhibitor, pepstatin A (PepA)(36). CatD localises to late endosomal/lysosomal compartments, whereas CatE has been found in a distinct set of endosomes of APCs; both have been implicated in antigen processing using inhibitors and knockout mice (34,37–42). Even though none of these studies reported any effects on MHCII expression levels, they did not rule out a role of CatD and/or CatE in MHCII protein turnover. To address this possibility, we here examined the accumulation of MHCII molecules following PepA treatment of various APC types; in addition, in a myeloid model APC line, we identified CatD as the enzyme responsible for the PepA effect and explored which forms of HLA-DR class II molecules are subject to proteolytic attack.

## Results

### PepA-mediated rescue of MHCII molecules in cultured APCs

To examine the role of aspartyl proteases in HLA-DR protein turnover, APCs were incubated with the aspartyl protease inhibitor, PepA (20 μM) for up to three days. Total (surface plus intracellular) amounts of mature (Ii-non-associated, heterodimeric) DR molecules were quantified by direct immunofluorescent staining of fixed, permeabilized cells with the monomorphic anti-DRα mAb, L243 (see Methods), followed by flow cytometry (**Fig. 1** and **Supporting Information, Fig. S1, A-C**). In the DR^+^ myeloid leukemic cell line, KG-1, HLA-DR accumulated following PepA treatment (compared with vehicle control) with approximately linear kinetics at ≈ 25% per day over the first 48 hours of culture (**Fig. 1A, B**). The accumulation seen in Fig. 1B at 48 hours was reproducible (mean 48% ± 28% increase compared to vehicle control) in multiple experiments (**Fig. 1C**). No accumulation was seen at the plasma membrane (**Fig. 1D**), implying that the rescued DR molecules were intracellular. Together, these findings implicated aspartyl proteases in HLA-DR protein turnover.

**Fig 1.**
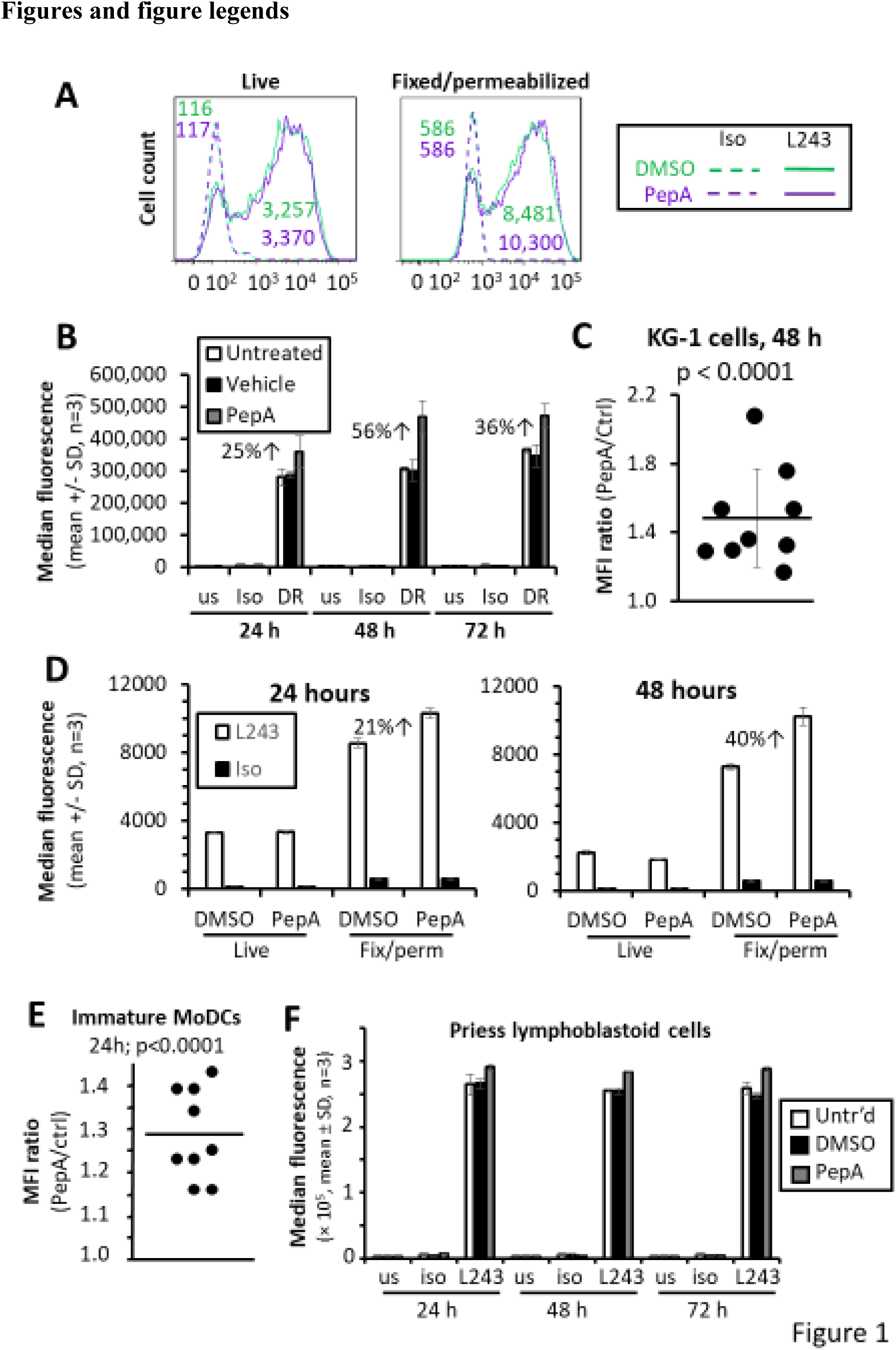
Rescue of HLA-DR molecules by PepA. A, Representative immunofluorescent anti-HLA-DR and isotype control staining profiles of live or fixed, permeabilized KG-1 cells treated with PepA (20 μM) or DMSO (vehicle control) for 48 hours. Flow cytometric analysis using FACSCantoII after gating for singlet intact cells. The gating strategy and similar data obtained using an Accuri C6 flow cytometer are shown in **Fig. S1**, **A-C**. B, Median fluorescence intensities (mean ± SD, n = 3 biological replicats) from a representative experiment as in (A) with daily sampling over 72 hours. Data from untreated and vehicle-treated controls are included, as well as unstained and isotype controls. Analysis using Accuri C6. A 2-way ANOVA of DR staining data indicated significant effects of time in culture (2 df, F = 27.5, p < 0.001) and treatment (2 df, F= 150, p < 0.001) and a significant time × treatment interaction (4 df, F = 4.2, p=0.013), consistent with a time-dependent effect of PepA. By Tukey’s HSD post-test, the PepA condition was significantly different from untreated and vehicle control (p< 0.001 for both). The staining on each day of analysis was done separately, so technical variation in staining may contribute to the time effect. C, Anti-DR staining ratios (L243-FITC MFI) of PepA-treated cells, divided by vehicle control) after 48 hours of culture, summarizing 9 independent experiments as in (B). P value was determined by 1-sample t test against an expectation value of 1.0 (representing no PepA effect). D, HLA-DR or isotype control staining of KG-1 cells after 24 h (left) or 48 h (right) of culture with PepA or DMSO control, comparing staining of live and fixed, permeabilized cells. At both time points, a 2-way ANOVA of DR staining data showed significant effects (p < 0.0001 each) of inhibitor treatment, of fixation and permeabilization, and of the interaction between these two factors. E, Immature MoDCs from healthy donors were differentiated for 7 days, then cultured with 20 μM PepA or vehicle control for a further 24 hours before analysis similar to (C), except that single measurements were performed (see **Fig. S3**). Shown are anti-DR staining ratios (L243-FITC MFI in PepA-treated cells divided by vehicle controls) from 9 independent experiments with MoDCs from 5 donors. F, Experiment as in Fig. 1B, but using the lymphoblastoid cell line, Priess. The small treatment effect in this attempt (≈ 10% increase) was statistically significant (2 df, F = 30.8, p < 0.001), but there was no statistical evidence for accumulation over time (4 df, p = 0.952 for interaction), and the increased anti-DR staining in PepA-treated cells was not reproduced consistently in 4 independent repeats.

Normal growth media may contain varying amounts of the lysosomotropic agent, ammonia, due to spontaneous hydrolysis of L-glutamine, resulting in a slow-down of lysosomal degradation (43, 44). To examine whether the PepA effect was observable in the absence of this potential confounding variable, KG-1 cells were switched to media containing GlutaMax, a non-hydrolyzable source of glutamine, and the experiment in Fig. 1B repeated (**Fig. S2A**). Again, PepA caused HLA-DR accumulation, to a comparable extent (32% maximal increase), but this time the effect peaked at 6-24 hours. Thus, the PepA effect was observed regardless of the source of glutamine and its effect on lysosomal activity, but the time of maximal PepA effect was affected by the source of glutamine, likely reflecting differences in lysosomal activity. Interestingly, DR accumulation levelled off after 48 hours in normal culture media (Fig. 1B) and subsequently diminished in GlutaMax media (**Fig. S2A**; cf. Discussion).

Because 20 μM was the highest dose of PepA achievable without toxicity, we examined whether an analogue engineered for increased intracellular bioavailability, PepA-penetratin, might achieve greater rescue of HLA-DR molecules. However, the rescue achieved with 10 μM PepA-penetratin was not substantially greater than with 20 μM unconjugated PepA (**Fig. S2B**).

As a specificity control, we previously reported that HLA class I molecules are not detectably rescued by PepA (or by any other protease inhibitors) in KG-1 cells (45). This suggests that CatD has no role in HLA class I degradation in this cell line, or that this enzyme acts redundantly with other lysosomal cathepsins on class I molecules.

In immature monocyte-derived dendritic cells (MoDCs, Fig. 1E and **Fig. S3**), PepA rescue of HLA-DR was similar in magnitude (mean ≈ 25% increased L243 staining after 24 hours) as in KG-1 cells at the same time point. We and others have shown previously that immature MoDCs exhibited HLA-DR turnover with half-lives of ≈ 1 day (9, 45)^2^, so that the 25%/d rate of DR accumulation due to PepA accounted for ca. half of the reported turnover rate. In the EBV-transformed B-lymphoblastoid cell line, Priess, a small amount of HLA-DR accumulation was observed in the presence of PepA in some experiments (Fig. 1F), but not others. This was consistent with the very slow HLA-DR protein turnover in Priess cells detected by ^2^H_2_O labelling (t_1/2_ > 60 h)^3^. Taken together, the data suggest that the PepA effects correlate with rates of HLA-DR protein turnover, where known.

The PepA effect was not confined to human APCs. We observed varying extents of rescue by PepA of several murine MHC class II proteins in splenic B cells and DCs (**Fig. S4**). Thus, the observed role of aspartyl proteases in HLA-DR protein turnover may extend to other MHCII molecules.

### Role of CatD in the degradation of HLA-DR proteins in KG-1 cells

Next, we sought to identify the aspartyl protease responsible for the effect of PepA on HLA-DR levels. The two lysosomal aspartyl proteases, CatD and CatE, share homology (46) but can be distinguished on Western blot by immunoreactivity and the fact that CatE, unlike CatD, exists as a disulfide-bonded homodimer (**Fig. 2**, A and **B**). KG-1 cells showed substantial expression of CatD, whereas Priess cells expressed much lower quantities of this enzyme, visible only on long exposure of blots, with a different processing pattern (Fig. 2A). This was consistent with the lack of PepA effect in Priess cells (cf. Fig. 1). Neither cell line expressed detectable CatE protein (Fig. 2B). Quantitative real-time RT-PCR confirmed the presence of messenger RNA for CatD, but not CatE (**Fig. S5**), and CatD expression in KG-1 cells was confirmed by flow cytometry (**Fig. S1C**). CatD and HLA-DR immunoreactivity are known to co-localize in late endosomal/lysosomal compartments (47, 48). Thus, CatD was the only expressed aspartyl protease that could plausibly account for the PepA effect.

**Fig 2.**
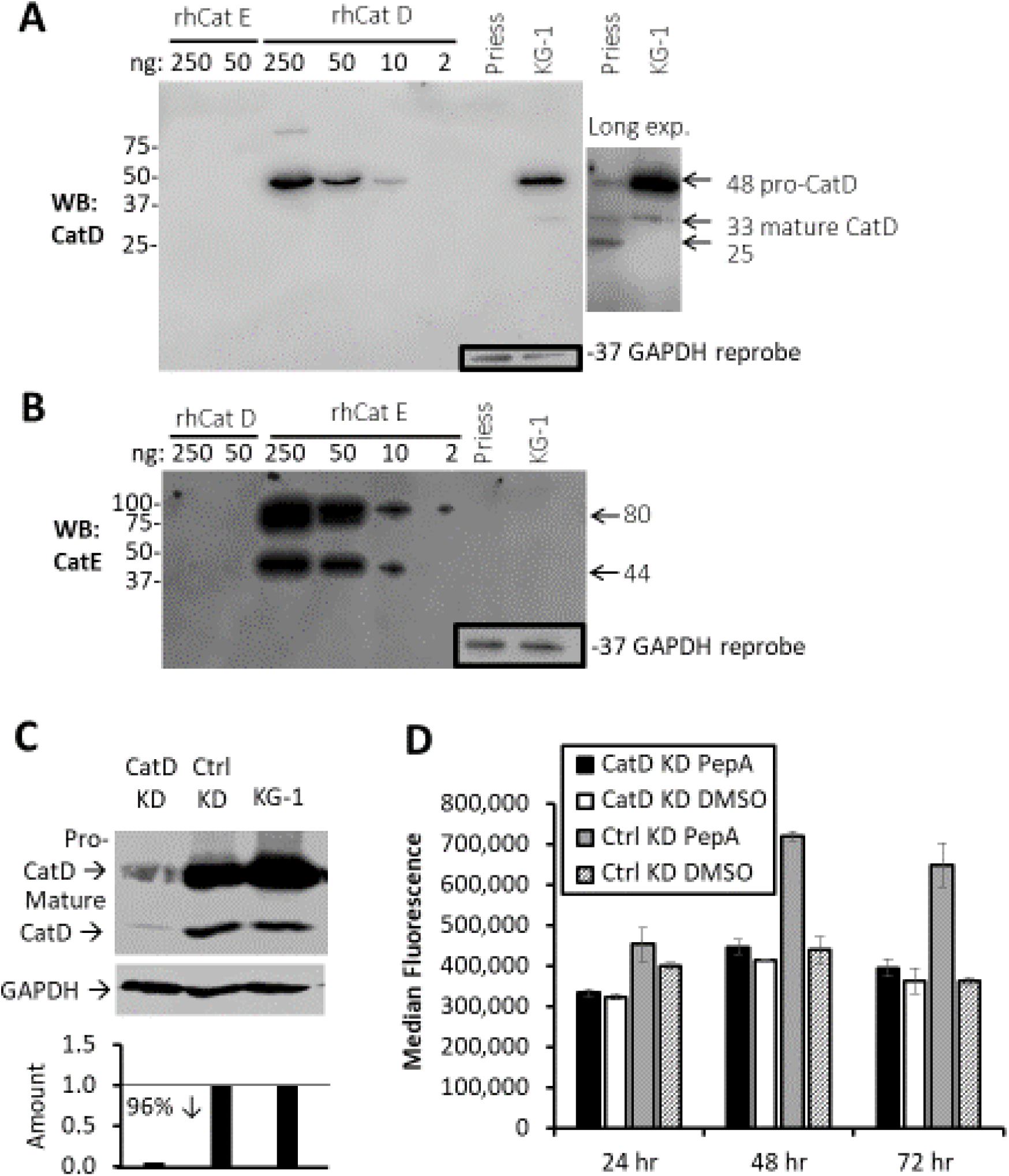
Role of CatD in HLA-DR degradation in KG-1 cells. A and B, Western blot analysis of KG-1 and Priess extracts using anti-CatD (A) and anti-CatE (B) antibodies. Titrated amounts of recombinant human CatD and CatE are included as specificity controls and semiquantitative standards. Insets show GAPDH loading controls (vertically displaced) for the Priess and KG-1 extracts after re-probing. C, shRNA-mediated knockdown of CatD protein expression, shown by anti-CatD Western blot of KG-1 cells lentivirally transduced with CatD shRNA knockdown construct (KD), compared to empty-vector control (*top*). After re-probing for GAPDH as a loading control (*middle*), band intensities were quantified, showing 96% knockdown (*bottom*). D, PepA effect on HLA-DR expression in CatD-knockdown and empty vector control-transduced KG-1 cells, analysed over 3 days. Means ± SDs of triplicates are shown. Analysis by 2-way ANOVA for PepA and knockdown effects at each time point. In the control knockdown cells, the PepA effect (1 df) was statistically significant at each time point (24 h: F = 81.2, p = 0.001; 48 h: F = 201, p < 0.001; 72 h: F = 81.0, p = 0.001). In CatD knockdown cells, the effect of PepA was not statistically significant at any time point (24 h: F = 9.6, p = 0.086; 48 h: F = 5.7, p = 0.05; 72 h: F = 0.1, p = 0.221)A and B, Western blot analysis of KG-1 and Priess extracts using anti-CatD (A) and anti-CatE (B) antibodies. Titrated amounts of recombinant human CatD and CatE are included as specificity controls and semiquantitative standards. Insets show GAPDH loading controls (vertically displaced) for the Priess and KG-1 extracts after re-probing. C, shRNA-mediated knockdown of CatD protein expression, shown by anti-CatD Western blot of KG-1 cells lentivirally transduced with CatD shRNA knockdown construct (KD), compared to empty-vector control (*top*). After re-probing for GAPDH as a loading control (*middle*), band intensities were quantified, showing 96% knockdown (*bottom*). D, PepA effect on HLA-DR expression in CatD-knockdown and empty vector control-transduced KG-1 cells, analysed over 3 days. Means ± SDs of triplicates are shown. Analysis by 2-way ANOVA for PepA and knockdown effects at each time point. In the control knockdown cells, the PepA effect (1 df) was statistically significant at each time point (24 h: F = 81.2, p = 0.001; 48 h: F = 201, p < 0.001; 72 h: F = 81.0, p = 0.001). In CatD knockdown cells, the effect of PepA was not statistically significant at any time point (24 h: F = 9.6, p = 0.086; 48 h: F = 5.7, p = 0.05; 72 h: F = 0.1, p = 0.221)

To test formally whether CatD was responsible for the PepA effect on DR expression, lentiviral shRNA knockdown of CatD was performed in KG-1 cells, yielding a 96% reduction in CatD expression (Fig. 2C). KG-1 cells transduced with empty vector maintained the effect of PepA on HLA-DR expression (Fig. 2D), with similar kinetics as seen before (cf. Fig. 1B), but CatD knockdown resulted in almost complete ablation of the PepA effect. This provided genetic evidence that the PepA effect, and by implication the turnover of HLA-DR in the wild-type cells, was dependent on CatD expression. Interestingly, steady-state expression of DR in the CatD-depleted KG-1 cells was not elevated above wild-type (Fig. 2D; cf. Discussion).

### CatD performs HLA-DR cleavage at αPhe54 *in vitro*

In order to examine whether CatD directly acts on HLA-DR ectodomains *in vitro*, we subjected purified recombinant soluble DR0402 to digestion by human CatD under mildly acidic (endosome-like) conditions and analysed cleavage products by SDS-PAGE and silver staining (Fig. 3). Little evidence of CatE cleavage was found, and CatD cleaved only a small proportion of offered DR molecules under these conditions (Fig. 3A, cf. Discussion). Nonetheless, three specific CatD cleavage fragments were detected by silver staining, which were not present in control reactions without enzyme, or when the enzyme was inhibited by PepA. The bands were excised and analysed by tandem mass spectrometry. The known cleavage preferences of CatD (after large hydrophobic residues) and trypsin (after Arg/Lys) are non-overlapping, so we compared tryptic digests of the fragments, looking for novel cleavage products in the CatD fragments that were absent from the intact DR bands (Fig. 3B**; Table S1** and **Fig. S6**). By mass spectrometric analysis, the largest CatD fragment (D1, 18 kDa) showed evidence of α-chain cleavage at the Phe54-Glu55 peptide bond, which is compatible with the known preference of CatD for cleavage after Phe or Leu. Moreover, we scanned the DRα amino acid sequence against the known cleavage preferences of CatD, considering the frequencies of all 20 amino acids at each of the four residues on either side of known CatD cleavage sites (cf. Experimental Procedures, **Tables S2-S4** and supporting text). According to this analysis, αPhe54 was the top-ranked cleavage site (Fig. 3C). Inspection of the HLA-DR crystal structure showed that αPhe54 was in the extended strand C-terminal to the α-chain helix, facing the bound peptide (Fig. 3D, shown in magenta)(26). This result is compatible with a model whereby CatD initiates direct attack on HLA-DR proteins, cleaving at αPhe54 (see Discussion). The smaller bands (D2, 16 kDa and D3, 14 kDa) contained α-chain peptides consistent with one or more secondary CatD cleavages: residue αPhe51 was close to αPhe54, whereas αLeu92was buried at the α2/β2 domain interface, distant from the peptide-binding groove (Fig. 3D, shown in light green at *right*; **Table S1**). Although tryptic β-chain fragments were observed (**Table S1** and **Fig. S6A**), they spanned most of the sequence, and of the few novel peptides observed, none matched the cleavage preferences of CatD (**Fig. S6B**).

**Fig 3.**
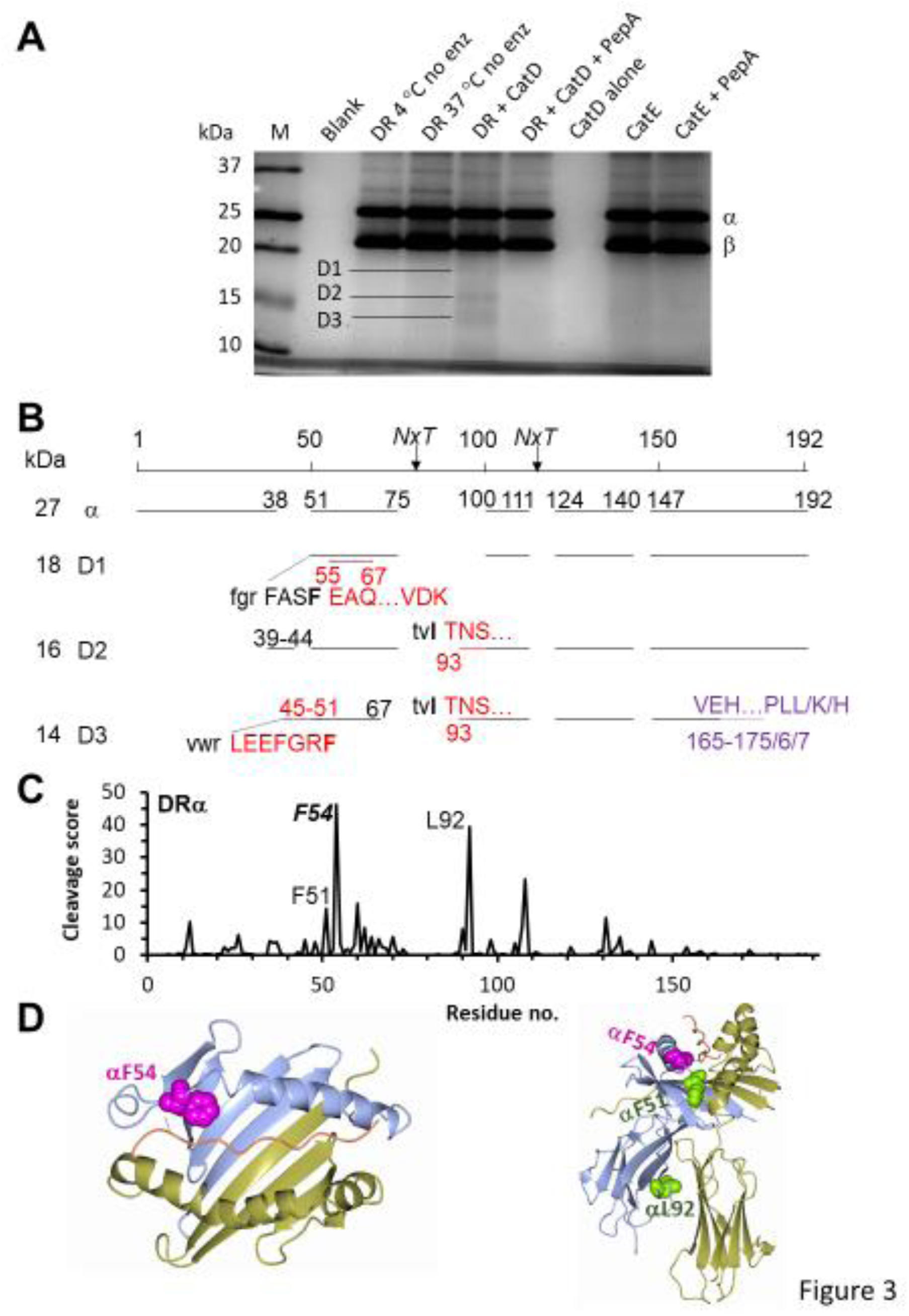
Cleavage of HLA-DR by CatD in vitro. A, Analysis of CatD digests. Purified soluble recombinant HLA-DR0402 protein was digested for 16 hours with or without CatD at 37 °C and pH 4.5, as shown, or kept at 4 °C. As a further negative control, CatD was blocked with excess PepA before digestion. Digests were resolved by SDS-PAGE, and the intact α and β chains, as well as the indicated fragments, were excised for tandem mass spectrometry. B, Schematic map of fragments (horizontal lines) found in the intact α chain and the CatD fragments D1/2/3 shown in (A). Black lines and numbers show the mapping of tryptic fragments to the expected amino acid sequence. Red lines and uppercase red text and numbers show additional peptides, which were absent from the intact α chain and compatible with CatD cleavage (cf. (C)). Purple dashed lines, text and numbers show peptides in the D3 fragment that were absent from the intact chain but were not explained by CatD cleavage. For analysis of β-chain-derived sequences, see **Fig. S6A**. The sequence alignments for both chains are shown in **Table S1**. C, Predicted CatD cleavage preferences in the HLA-DR α chain. The Y axis indicates relative predicted cleavage probability (Experimental Procedures, **Tables S2**-**S4** and supporting text); the Y axis indicates primary sequence position, aligned with the map in (B). The novel observed fragments are compatible with peak cleavage probabilities. D, Views of the DR0101-CLIP crystal structure (PDB: 3QXA) showing the location of the residues in the P1 position relative to the observed CatD cleavage sites. Polypeptide main chains are color-coded blue (DRα), gold (DRβ), pink (CLIP). Space-filling models of side chains are shown in magenta for the initial cleavage site at αF54, and in green for the secondary cleavage sites (αL92, αF51). *Left*, top view of the peptide-binding groove showing only the initial cleavage site. The main-chain nitrogen of αF54 hydrogen-bonds with CLIP (dashed line). *Right*, side view of the entire structure (with CLIP N terminus pointing towards the reader), with the locations of all cleavage sites shown.

### Characterisation of HLA-DR molecules rescued by PepA in KG-1 cells

HLA-DR associates with different molecules during its intracellular maturation and fate (see Introduction), so we examined which forms of DR were rescued in KG-1 cells after 2 days of culture with PepA. First, noting that DR molecules loaded with stable peptides acquire resistance to SDS denaturation at room temperature (49–51), SDS stability was evaluated by Western blot (Fig. 4A). Antibodies to the DR α or β chain were used, which had different preferences for SDS-unstable vs. SDS-stable forms of DR. Regardless of this, the band intensity of SDS-stable αβ/peptide heterotrimers was unaffected by PepA, whereas the abundance of SDS-unstable molecules almost doubled in extracts from PepA-treated vs. control KG-1 cells (Fig. 4B, *left*). When the extracts were boiled, the SDS-stable bands disappeared and the intensity of the monomer bands increased, as expected (Fig. 4A); quantification showed an increase in PepA-treated cells comparable to that seen by flow cytometry (cf. Fig. 4B, right and Fig. 1B). Together, these findings implied that PepA did not rescue DR molecules that acquired stably bound peptide and focussed further analysis on other forms of DR that are not SDS-stable.

Importantly, the DR molecules rescued by PepA were not detectably ubiquitinated, which would have produced a ladder above the main β-chain monomer band with an 8 kDa spacing (Fig. 4A, *right*). Neither did any lower-molecular weight fragments accumulate in the presence of PepA, suggesting that intact molecules, rather than partial cleavage products generated by other enzymes, were rescued.

**Fig 4.**
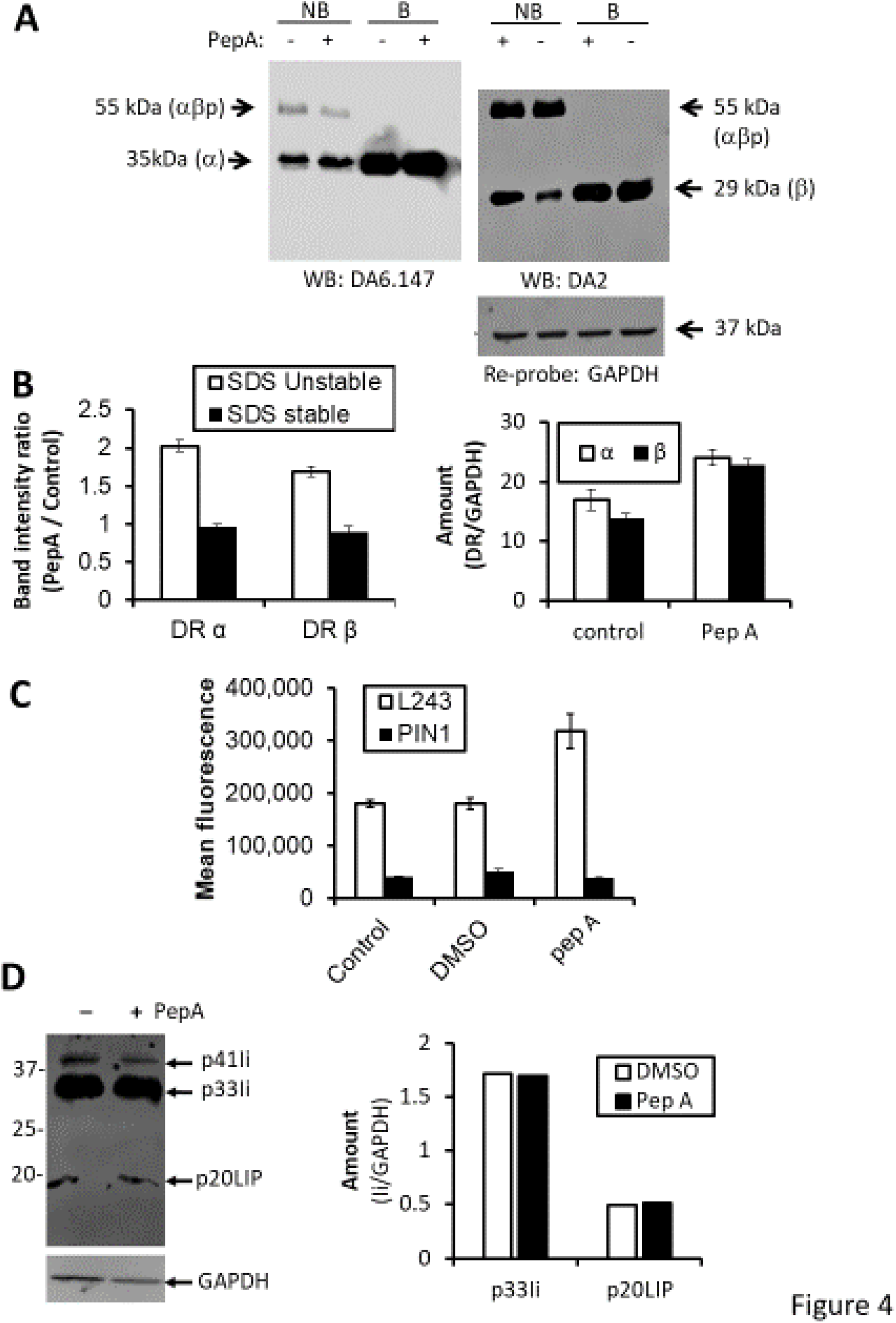
Characterisation of the forms of HLA-DR rescued by PepA. A, SDS stability. KG-1 cells were grown for 2 days in the presence of 20 μM PepA or vehicle control (DMSO) as indicated. Total protein-matched, non-boiled (NB) or boiled (B) cell extracts were resolved by 12% nonreducing SDS-PAGE, and Western blots probed with antibodies to HLA-DR α (*left*) and β chains (*right*). The blot for the β-chain was re-probed for GAPDH (*bottom right*). B, Quantification of band intensities in A. *Left*, band intensities representing monomers and SDS αβ/peptide complexes in non-boiled samples from A were quantified by using LiCor imager, and the effects of PepA shown as a bar graph. *Right*, GAPDH-normalised band intensities in boiled samples were used to quantify rescue of total DR molecules. Bar graphs show mean ± SD of two independent experiments. C, Flow cytometric analysis for Ii. KG-1 cells were treated with or without PepA as in B, fixed and permeabilized, and analysed by flow cytometry after staining with a mAb specific for the cytoplasmic tail of Ii (PIN.1) or, as a positive control, with mAb against mature, Ii-non-associated DR (L243). An example of staining with specificity controls is shown in **Fig. S1D**. At 48 h, the anti-DR staining differed significantly between culture conditions (1-way ANOVA, 2 df, F = 18.2, p < 0.001), but the anti-Ii staining did not (2 df, F = 1.4, p = 0.28). D, Western blot analysis for Ii. Extracts from KG-1 cells treated for 48 hours with or without PepA were resolved by SDS-Page and Western blots probed with anti-Ii mAb PIN.1 (*left*). The intensities of the p33Ii and p20LIP bands were quantified (*right*). The experiment was performed only once, but the results conformed to those in Fig. 4D and Fig. S2A.

The mAb used to detect DR αβ dimers in Fig. 1, L243, reacts poorly with Ii-associated DR (cf. Methods), suggesting that the L243-reactive DR molecules rescued by PepA were not Ii-associated. Nonetheless, we examined this possibility directly, because previous studies had provided evidence for (52) and against (40,53,54) aspartyl protease involvement in the initial processing of the class II-associated Ii; variation in this regard may be due to differences in lysosomal protease composition or HLA class II genotype (55). In KG-1 cells, however, PepA failed to mediate any detectable rescue of Ii or its fragments, as shown both by flow cytometric analysis with Ii-specific mAb (Fig. 4C and **Fig. S1E**; time course in media with glutamine or GlutaMax shown in **Fig. S2A**, *bottom*) or by Western blot analysis of full-length Ii and its degradation intermediates (Fig. 4D). Thus, the intracellular accumulation of DR in PepA-treated KG-1 cells was not due to arrest of Ii processing in the endocytic pathway. The terminal Ii processing product, DR-associated CLIP, was not detectable in the KG-1 cells by flow cytometry with CLIP-specific antibody (**Fig. S1E**); at least one of the two coexpressed DRB1 alleles in KG-1 cells, DRB1*11:01, has been reported to have only modest CLIP affinity (56).

Fig. 5A compares the biochemical characteristics of the PepA-rescued DR molecules with the known characteristics of the different maturation stages of DR. The findings are consistent with the rescue of empty or poorly-loaded DR molecules from degradation by PepA; in contrast, neither ubiquitylated DR molecules, SDS-stable complexes containing stably-bound peptides, or DR molecules associated with Ii or Ii degradation intermediates, such as p20LIP, were rescued by PepA.

**Fig 5.**
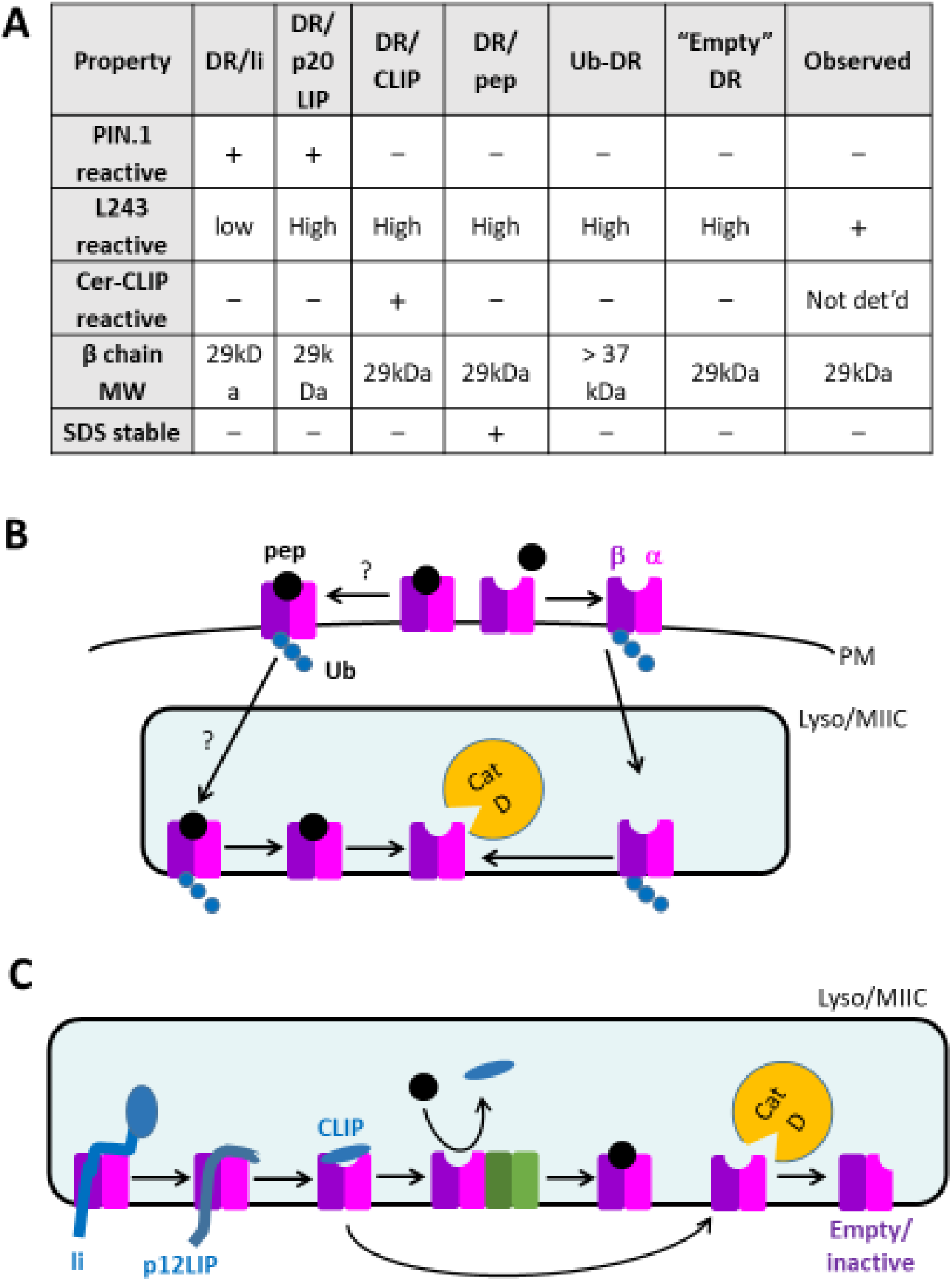
Working model of CatD-mediated HLA-DR protein turnover. See text for details and citations. A, Table comparing the biochemical characteristics of PepA-rescued DR molecules (from Fig. 4) with various stages in DR maturation. B, Postulated role of CatD in the initial proteolytic attack on empty MHCII molecules following β-chain ubiquitylation, internalization to lysosomes, de-ubiquitylation, and loss of bound peptide. Selective attack on empty (or poorly-loaded) MHCII molecules may occur because hydrogen-bonding with stably-bound peptide prevents access of CatD to the scissile bond at αF54 (cf. Fig. 3D); in addition, it is conceivable that empty/poorly-loaded DR may be preferentially ubiquitylated and internalized, as compared to DR molecules stably loaded with peptide (indicated by “?”). C, Molecular interactions with Ii, DM and peptide along the classical MHCII maturation and peptide loading pathway would prevent initial CatD attack on the scissile bond at αF54. Conformational changes of empty DR following peptide release may also influence CatD susceptibility.

## Discussion

This study identified a novel role of the aspartyl protease, CatD, in the degradation of MHCII proteins. MHCII expression level and lifespan have been linked to peptide loading (7,20,21) and are regulated in APCs by β-chain ubiquitylation and internalization (11, 12). Using the aspartyl protease inhibitor, PepA, expression analysis and shRNA knockdown, we showed in a well-characterised model APC line that CatD is involved in the degradation of HLA-DR molecules. By targeting a site on DR that is involved in “locking” peptides into the peptide-binding groove, CatD seems to target DR molecules that are not associated with Ii or stably bound peptide. This explains well-established links between accelerated MHCII protein turnover and impaired peptide loading. A model accounting for these findings is shown in Fig. 5B.

Mechanistic studies focused on the KG-1 model cell line, but the findings are likely to be more broadly applicable. Rescue of HLA-DR molecules by PepA was observed in immature MoDCs, a well-characterised APC type (57), which was previously shown to exhibit MHC class II protein turnover (9, 45)(Fig. 1E**, Fig. S3**). PepA effects were seen in MoDC from unrelated donors,^2^ suggesting that various DR alleles are similarly susceptible to aspartyl proteases. This is consistent with the finding that CatD was found to attack the non-polymorphic DRα chain *in vitro*. PepA was also able to rescue murine MHCII molecules in splenic APCs, albeit with weaker effects in CD11c+ DCs than in B cells and some allele-dependence (**Fig. S4**). Residue F54 is conserved in the amino acid sequences of human and mouse MHCII α chains, albeit with variation in flanking sequences that may affect CatD susceptibility, potentially confining CatD susceptibility at this site to HLA-DR, -DP and H2-E α chains (**Table S5**). H2-A^d^, which is CatD susceptible experimentally (**Fig. S4**), scores lower but has the highest score of the H2-A alleles at αF54; uniquely among H2-A alleles, it has a potential alternative cleavage site at αL53, immediately adjacent in the same strand (**Table S5**).

Of the two aspartyl proteases, CatD, which is responsible for the PepA rescue in the KG-1 model, co-localizes with MHCII in late endosomes and lysosomes, and it is therefore available to carry out MHCII protein degradation in any APC type (34,37,40).^4^ CatE shares sequence homology and sequence-dependent cleavage preferences with CatD (58) but, owing to its existence as a disulfide-bonded homodimer, may have more stringent steric requirements than the monomeric CatD; it is expressed in APCs but not found in late endosomes or lysosomes (39). In KG-1 cells, CatE expression was undetectable, and the PepA effect was clearly CatD-dependent, as shown by CatD knockdown. *In vitro*, evidence for CatE attack on recombinant soluble DR was weak. Nonetheless, we cannot exclude the possibility that CatE could contribute to MHCII degradation in APCs that express this enzyme.

Comparing the rate of DR turnover in MoDCs (*t1/2* ≈ 1 day, equivalent to 50% new DR protein per day) with the rate of DR accumulation in PepA-treated MoDCs (25% increase per day in standard media) suggested that aspartyl protease activity accounts for very approximately half of the observed rates of DR turnover in this system. The remainder could be accounted for by the existence of non-aspartyl proteases with redundant functions (see below) or by the loss of some MHCII molecules present on exosomes (59, 60). It is difficult, however, to rule out the alternative possibility that inhibition of CatD activity was incomplete at the maximal non-toxic dose of PepA used. In the lymphoblastoid cell line, Priess, lower CatD expression correlated with slower turnover^3^ and little if any DR rescue by PepA (Fig. 1F), compared with MoDCs, although other factors, such as ubiquitylation, may contribute to these differences (61).

Our data may help to explain how MHCII molecules survive proteolysis by CatD and other lysosomal enzymes during their maturation and peptide loading, yet are degraded in the same compartments. We propose that, during maturation, the site of initial CatD attack is protected by interactions with invariant chain, DM or bound peptide; in contrast, poor loading or loss of bound peptide may leave DR molecules vulnerable to CatD attack (Fig. 5C). Initial CatD attack occurs at residue αPhe54 (Fig. 3D), in an extended strand C-terminal to the α-chain helix, which hydrogen-bonds with CLIP and antigenic peptides in MHCII crystal structures (26,62,63). In addition, interaction with intact Ii prior to its degradation may sterically mask the vicinity of the CatD attack site by its trimerization domain, C-terminal to CLIP (64, 65). When DR binds to DM during peptide exchange (66), the extended strand around the site of CatD cleavage at αF54 changes conformation but remains sterically protected within the DM/DR complex. Thus, this site remains unavailable to CatD attack throughout all steps of the MHCII peptide loading pathway. These considerations may explain why, in KG-1 cells, DR molecules resisted CatD attack while associated with Ii, Ii fragments, or stably bound peptide. In contrast, other SDS-unstable forms of DR (likely DR molecules with loosely bound peptide, or none) were vulnerable to CatD attack (Fig. 4).

Molecular dynamics simulations of empty DR molecules, and biophysical studies of mutant DR molecules with altered peptide-exchange characteristics, suggest that the extended strand around αPhe54 gains conformational flexibility and, partly due to molecular movements in the opposing β-chain helix, becomes more accessible in peptide-receptive empty conformations (67–69). This would provide steric access for CatD to this part of the peptide-binding groove in empty molecules, explaining the selective degradation of such molecules. Intriguingly, αF54 is not only a key site of CatD attack, but is also implicated in these conformational changes by mutagenesis studies (69).

Only a small proportion of DR molecules could be degraded by CatD *in vitro* in our experiments. For other substrates, the optimal pH for CatD attack has been reported as pH2-3.5 (36, 70), but we explored only lysosomal pH values (4.5-5.5), in order to maintain DR folding (71) and thus physiological relevance. The insect cell-derived recombinant soluble DR molecules used in these studies are not loaded with an edited, SDS-stable peptide repertoire, unlike native DR molecules purified from mammalian APC. Nonetheless, their peptide-binding grooves have been found to be at least partially occupied by polypeptides (72). In addition, kinetic and molecular dynamics studies suggest that peptide-receptive conformations are metastable, collapsing into inactive conformations (73), in which the strand on either side of residue αF54 moves into the peptide-binding groove (69,74,75). Aggregation of empty MHCII molecules (76, 77) may also hinder CatD attack. Open, monomeric conformations that are vulnerable to CatD attack may therefore be rare. Taken together, these factors may help to explain why only a small percentage of DR molecules was degraded by recombinant CatD *in vitro*, and they may contribute to the slow rate of DR turnover *in vivo*. Although further experiments will be required to elucidate how peptide binding and conformational states influence the CatD susceptibility of HLA-DR, we hypothesise that initial CatD attack on HLA-DR at αF54 may selectively target an unstable, peptide-receptive conformation (Fig. 5C).

*In vitro*, CatD cleaved DR molecules not only after αF54, but also after other large hydrophobic residues, producing smaller fragments (Fig. 3D); however, one of these residues is buried in the folded structure of DR, and a third is near the initial cleavage site; cleavage at these residues may reflect secondary degradation steps. *In vivo*, the DR polypeptides rescued from degradation by PepA were intact molecules rather than smaller degradation intermediates; this is consistent with the idea that CatD is important for initiating proteolytic attack, although it may also contribute to the downstream degradation steps, alongside other proteases.

DR molecules could become available for CatD attack at different times during their post-translational fate. The first opportunity for degradation would arise from a failure to acquire stable peptides following Ii processing, upon release of CLIP and dissociation from DM (Fig. 5C). Indeed, in cell lines with defective peptide loading due to DM deficiency, pulse-chase analysis shows some attrition of DR molecules relatively shortly after biosynthesis (24). However, the much longer half-life of bulk HLA-DR molecules and the slow kinetics of the PepA effect suggest that this is not the dominant pathway in the APC that we have studied. A subsequent opportunity for CatD attack would arise when, after a time of residence at the plasma membrane, DR molecules are ubiquitylated and re-internalised (14). We found no evidence that the DR molecules rescued by PepA in KG-1 cells remained ubiquitinated (which would have raised the molecular weight of the β-chain in PepA-treated cells); rather, any cytoplasmic oligo-ubiquitin chains may be removed by de-ubiquitinating enzymes prior to CatD attack on the ectodomains (78, 79). An unresolved question is whether the remarkable selectivity for SDS-unstable, Ii-non-associated DR molecules (Fig. 4) is exclusively due to the inherent specificity of CatD, or whether, in addition, ubiquitin-dependent internalisation selectively targets DR molecules that have lost bound peptide for lysosomal degradation (Fig. 5B).

Several findings indicated that KG-1 cells restore HLA-DR levels to wild-type after prolonged inhibition (**Fig. S2A**) or genetic ablation (Fig. 2D) of CatD. Conceivably, new HLA-DR synthesis could slow down to match the reduced rate of degradation. Alternatively, lysosomal overload caused by a lack of CatD activity could prompt the induction of other lysosomal enzymes that compensate for the defect. A possible precedent for the latter mechanism has been found, albeit in the opposite direction: CatD levels were shown to be elevated in the skin of CatL-deficient mice (80). Adaptation to a chronic lack of active CatD might help to explain why previous studies in CatD knockout mice (40) and genome-wide screens for modifiers of DR expression (32) have failed to detect the role of CatD in DR degradation identified here. In those studies, there would have been ample time for adaptive mechanisms to compensate for the lack of CatD. For the same reason, genetic studies of spontaneous or engineered defects in lysosomal proteases might underestimate their roles in normal biology. In human neuronal ceroid lipofuscinosis, a severe congenital lysosomal storage disorder attributed to hypomorphic mutations in CTSD (CatD gene; OMIM entry 116840), the neurodevelopmental consequences of CatD insufficiency dominate the clinical manifestations, and any immune abnormalities may be comparatively minor or masked by adaptation, or both.

MHCII allelic variation is a major determinant of autoimmune disease risk (22,81–83). Instability of the disease-associated MHCII alleles has been proposed as a mechanism for impaired tolerance in NOD mice and human T1D patients with HLA-DQ risk alleles (7). Recent studies from our laboratory, however, using ^2^H_2_O labelling techniques to quantify turnover in NOD APCs *in vivo*, found no evidence for an intrinsic stability defect of the risk alleles, and no correlation with autoimmunity (7, 8). The present study shows that CatD attack on HLA-DR involves a monomorphic cleavage site (Fig. 3), and our unpublished studies show little variation in HLA-DR half-lives between individuals.^2^ Both findings argue against a major role for allelic differences in DR protein turnover in HLA-DRB1-linked risk of autoimmunity. Moreover, if our findings in KG-1 cells are representative of physiological APCs, blocking of DR protein degradation by CatD would not restore any presentation defect, as the rescued DR molecules accumulate intracellularly, and thus would not contribute to antigen presentation even if they were weakly bound to self-peptides.

### Concluding remarks

We found that CatD initiates HLA-DR degradation by selective cleavage at a site whose susceptibility to attack is affected by interactions with Ii, DM and peptide loading. CatD acts selectively on DR molecules lacking bound Ii or stably bound peptide, providing a means of lysosomal quality control. The data explain how HLA-DR molecules resist degradation during their maturation but are degraded in APCs as needed. They argue against a role of allelic differences in HLA-DR protein instability as a mechanism of DRB1 gene associations with autoimmunity. Our work raises novel questions about the molecular mechanisms linking HLA-DR peptide-loading state to internalization vs. degradation. Adaptive mechanisms may exist that restore wild-type DR levels in cells lacking CatD activity. The ability to trap HLA-DR molecules that would otherwise be degraded will enable further dissection of this immunologically important disposal pathway.

## Experimental Procedures

### Cell culture and inhibitor studies

Priess is a DRB1*04:01-homozygous lymphoblastoid cell line (84). KG-1 is an acute myleloid leukemia cell line (85) with features resembling immature DCs (86, 87), expressing DRB1*11 and *14 (88). Cells were grown at 37 °C in a humidified incubator, in RPMI1640 medium supplemented with 10% FBS, 2 mM L-glutamine, and penicillin/streptomycin, except for KG-1 cells, which were grown in IMDM with 20% FBS, penicillin/streptomycin, glutamine, and 25 mM HEPES. In some experiments, 2 mM L-glutamine was replaced by 1% v/v GlutaMax (L-alanyl-L-glutamine)(89). For inhibitor studies, triplicate cultures were set up with 20 μM PepA (diluted from 5 mM stock in DMSO), with an equivalent volume of DMSO as a vehicle control. These conditions did not interfere with the growth kinetics (by hemocytometer counting) or scatter profile (by flow cytometry) of KG-1 cells. Alternatively, cells were treated with 10 μM Pepstatin A-penetratin (diluted from 1 mM stock in PBS), a membrane-permeant cationic peptide conjugate of PepA with greater intracellular bioavailability (90). Cells were harvested up to 72 hours thereafter for analysis.

### Lentiviral shRNA knockdown

A commercial MISSION CTSD shRNA construct in pLKO.1-puro vector (MISSION, Sigma-Aldrich) was used for CatD knockdown; empty vector was used as a negative control. KG-1 cells (2.5 × 10^5^ in 200 μl media) were treated with 8 μg/ml polybrene overnight in a flat-bottom 96-well plate. Lentiviral supernatants were added at MOI 5 and the plate centrifuged at room temperature for 30 min. at 500 × *g*. Cells were expanded into larger plates, selected with 1 μg/ml puromycin, and expanded further into culture flasks for experiments.

### Monocyte-derived dendritic cell culture

Monocyte-derived dendritic cells were derived from healthy donor blood as described (45) with modifications as follows. Briefly, volunteers were recruited with ethical approval (Cambridge Research Ethics Committee 01/363) and informed consent in accordance with the Declaration of Helsinki. Heparinised venous blood (10 ml) was withdrawn, and mononuclear cells were isolated over Ficoll gradients. CD14^+^ monocytes were immunomagnetically enriched using Miltenyi CD14 microbeads and cultured for 6 days in complete RPMI media with GM-CSF and IL-4, yielding CD11c^+^ CD14^-^ MoDCs that were phenotypically immature (CD86^low^, HLA-DR^low^) as reported earlier (supplemental data in (45)). They were treated with 20 μM PepA or DMSO vehicle for another 24 hours. The LPS antagonist, polymyxin B (20 μg/ml)(91), was added 1 h before inhibitor treatment to minimize any inadvertent LPS activation.

### Quantitative real-time RT-PCR analysis

RNA was isolated from 10^5^ KG-1 or Priess cells, reverse-transcribed and PCR-amplified using the Power SYBR Green Cells-to-Ct Kit (Life Technologies). Quantitative real-time PCR was performed in 96-well plates using the SYBR Green method and a Step One Plus qPCR instrument (Applied Biosystems). After 10 min. at 95 °C for enzyme activation, 30 cycles of denaturation (95 °C, 15 s), annealing (55-60 °C optimised for each primer set, 1 min.), and extension (72 °C, 15 s) were performed, followed by a meltcurve at a heating rate of 1 °C per min. Primers for CatD, CatE, and GAPDH (housekeeping control) were as described (92–94); predicted amplicon sizes were, respectively, 296, 500 and 206 bp. PCR products were resolved by 0.7% agarose gel electrophoresis in TBE (70V, 90 min.), alongside a 100-bp DNA ladder (Promega), and visualized using SYBR Safe dye (Invitrogen) and a LiCor Odyssey Fc imager.

### Immunofluorescent staining and flow cytometry

For surface staining, human cells were suspended in phosphate-buffered saline (PBS) with 1% BSA and 0.05% sodium azide (FACS buffer), blocked for 15 min. with 10% normal mouse serum, and incubated with saturating concentrations of anti-HLA-DR mAb-FITC conjugate (BioLegend)(95) for 30-60 minutes. Cells were washed in FACS buffer and analysed immediately or, after washing in PBS, fixed in Cytofix (BD) for later analysis. The mAb clone used, L243, reacts poorly, if at all, with MHCII αβ dimers that are associated with full-length Ii but well with subsequent maturation stages (49,96–99).

For total (surface plus intracellular) DR staining, cells were washed in PBS, fixed and permeabilized using the Cytofix/Cytoperm kit (BD), washed in 1× Perm/Wash buffer, and subsequently processed as for live staining, except that all reagents were diluted, and cells washed, in 1× Perm/Wash buffer. PIN.1, an IgG1 mAb to the human Ii cytoplasmic tail (Santa Cruz Biotechnologies)(99), was used to detect Ii. CatD was detected by indirect staining, using primary polyclonal rabbit anti-CatD IgG (Abcam; ab75852) and secondary APC-conjugated anti-rabbit IgG. Unstained, secondary antibody alone, and isotype control stains were included as appropriate.

Flow cytometric analysis was performed using either FACSCantoII or Accuri C6 flow cytometers (BD). Data were acquired for at least 5000 cells per sample, with gating for intact singlet cells, and median fluorescence values were determined.

### Analysis of murine splenocytes

Animal studies were performed in accordance with institutional and UK Home Office ethical and licencing requirements (Project Licence 80/2156). Pre-diabetic nonobese diabetic (NOD, expressing H-2A^g7^) and Balb/c (H-2A^d^, E^d^) mice were crossed and F1 offspring aged to 6 weeks (8). After CO2 asphyxiation and cervical dislocation, spleens were removed and splenocyte suspensions obtained aseptically by Liberase CI (Roche) digestion, according to manufacturer’s instructions, followed by passage through a 70 μm mesh. Splenocytes were cultured in complete RPMI for 24 hours with 20 μM PepA or vehicle as described above. For flow cytometric analysis, cells were stained for DC (CD11c) and B-cell (B220) lineage antigens and, after fixation and permeabilization, with mAbs specific for each of the three MHCII molecules (8).

### Western blot

Using modifications of previously-described protocols (49), cell pellets were extracted in RIPA buffer (100) with Roche Complete protease inhibitors, normalised for protein content using bicinchoninic acid (BCA) protein assay, mixed with SDS-PAGE sample buffer, boiled for 5 minutes, and run on 12% SDS-PAGE mini-gels (Bio-Rad). For SDS stability assays, the boiling step was omitted where indicated. Resolved proteins were transferred to PVDF membranes, which were blocked with 5% nonfat dry milk in PBS, probed with antibodies to monomorphic determinants of HLA-DR α and β chains (mAbs DA6.147 and DA2, respectively, both from Santa Cruz Biotechnologies)(101, 102), invariant chain (mAb PIN.1, see above), CatD (mAb 185111, R&D Systems) or CatE (polyclonal goat IgG, Cat. No. AF1294, R&D Systems). Some blots were stripped and re-probed with mAb against GAPDH (Santa Cruz Biotechnologies) as a loading control. Bound antibody was visualised using appropriate HRP-conjugated second-step reagents, chemiluminescent detection, and LiCOR Odyssey Fc imaging and band quantification.

### *In vitro* digestion

Recombinant soluble HLA-DR0402 proteins (ectodomains encoded by DRA and DRB1*04:02) (103) were produced in insect cells and purified from culture supernatants by L243-sepharose immunoaffinity chromatography as described (71). Recombinant human prepro-cathepsins D and E (R&D Systems) were preincubated for auto-activation as per manufacturer’s recommendations (e.g., for CatD, 20 μg/ml enzyme in 0.1 M sodium acetate buffer, 0.2 M NaCl, pH 3.5; 30 min. at 37 °C). Purified DR0402 (4 μg/3 μl PBS/reaction) was digested with 4ng/2μl activated CatD or E (or buffer control), diluted to 20 ul in assay buffer (final pH 4.5). Where indicated, excess PepA was added before the enzyme (7 nmol/1 μl ethanol). Samples were incubated at 37 °C for 16 hours, neutralized with 1.5 M Tris-HCl, pH 8.8. Samples were resolved by SDS-PAGE and proteins visualized by silver staining. Bands were excised for LC/MS analysis.

### Mass spectrometry

Bands of interest were subjected to in-gel tryptic digestion and analysed by UPLC-MS/MS on an Orbitrap Velos mass spectrometer as previously described (45, 104). For the identification of CatD cleavage products, we exploited the orthogonal cleavage specificities of trypsin (after Lys/Arg) and CatD (after large hydrophobic residues (58, 105) (www.ebi.ac.uk/MEROPS_database entry A01.009 (106)). High-resolution (30, 000) Orbitrap Velos LC-MS/MS data files for tryptic digests of the intact DR α and β chains or specific cleavage products were screened, using MASCOT (Matrix Science), for matches to the sequences of the sDR0402 α and β chains (103), and the identified fragments were aligned to the DRA*01:01 and DRB1*04:02 ectodomain protein sequences from the IMGT/HLA database (https://www.ebi.ac.uk/ipd/imgt/hla/)(107). In addition to tryptic fragments, we screened for novel fragments that were not observed in the parent polypeptides, representing CatD cleavage.

### Protein structure analysis and visualization

Human HLA-DR, -DP and -DQ α-chain sequences were obtained from the IMGT HLA database. Murine H2-A and -E α-chain sequences were from the NCBI protein database (https://ncbi.nlm.nih.gov/protein). Protein structures were from the RCSB Protein Data Bank (https://www.rcsb.org/). Alignment of sequences (residues 41-70, mature DRα protein sequence numbering) was performed after inspection of crystal structures of DR, DQ and H-2A complexed with CLIP (PDB accession numbers 3QXA; 1A6A; 5KSU; 1MUJ). Putative CatD cleavage sites were visualized in the crystal structure of DR1-CLIP (3QXA) using CCP4MG software (http://www.ccp4.ac.uk/MG/)(108).

### Prediction of CatD cleavage from sequence

To check the plausibility of the experimentally identified cleavages, the observed CatD cleavage sites were compared with sequence-dependent predicted CatD cleavage probabilities, calculated in Microsoft Excel using input from the MEROPS database (106). Briefly, the frequencies of the 20 amino acids within an 8-residue window (P4 to P4’) around the cleavage site were obtained from the MEROPS specificity matrix for CatD. They were converted to probabilities and scaled by dividing by 5% (expectation value for 1 of 20 amino acids) to obtain scores for their over- or under-representation within CatD cleavage sites. For every potential cleavage site, the scores of the amino acid residues within the P4-to-P4’ sliding window were multiplied to obtain a cleavage likelihood score. An example of the computation is shown in **Tables S2-S4** and related supporting text.

### Data analysis

Flow cytometry data were processed in FlowJo or CSampler software. Band intensities on Western blots were quantified using LiCor Odyssey Fc ImageStudio software. Numerical data and descriptive statistics were processed in Microsoft Excel, and gel images exported into PowerPoint for annotation. Inferential statistics (cf. figure legends) were performed with IBM SPSS, R, or GraphPad Prism.

## Data availability

All data are contained within the manuscript.

## Acknowledgements

We thank the Phenotyping Hub of the Cambridge Biomedical Research Centre and the Department of Life Sciences, University of Roehampton, for access to flow cytometers. We are grateful for expert technical support and provision of critical reagents and services by staff in the Department of Life Sciences at the University of Roehampton; the Division of Human Gene Therapy, Department of Pediatrics at Stanford University Medical Center; and the Division of Rheumatology, Department of Medicine and the Cambridge Centre for Proteomics at the University of Cambridge. Dr Claudia Prevosto and Aleksandra Gumienny generated unpublished data mentioned in this paper. We thank Professor J.S. Hill Gaston for facilitating the recruitment of healthy human subjects and for critically reading the manuscript, and Dr Timo Burster for early discussions.

## Author contributions

Conceptualization: RB, SM Data curation: RB, NS, MJD Formal Analysis: RB, NS, ADR, MJD Funding acquisition: RB Investigation: NS, SM, CTAP, RWMR, AO, ADR, MJD, RB Methodology: RB, ADR Project administration: RB Resources: EDM, MJD Supervision: RB, MJD, EDM Visualization: NS, SM, CTAP, RWMR, AO, ADR, RB Writing – original draft: RB Writing – review & editing: all coauthors

## Funding and additional information

We are grateful for financial support from Versus Arthritis (formerly Arthritis Research UK; Senior Research Fellowship, ref. 18543, and Research Progression Award, ref. 20648), Diabetes UK (Small Grant), the MS Society (Project Grant, ref. 42), and the University of Roehampton (all to RB) and NIH AI095813 (to EDM). NS was supported by a MS International Federation McDonald fellowship for part of this work.

The content is solely the responsibility of the authors and does not necessarily represent the official views of the National Institutes of Health.

## Footnotes

1 Abbreviations used in this paper: APC, Antigen-presenting cell; CatD/E/G, Cathepsin D/E/G; CD, Cluster of Differentiation; CLIP, Class II-associated invariant chain peptides; HLA, Human leukocyte antigen; Ii, invariant chain; LIP, Leupeptin-induced polypeptide; MARCH-1, Membrane-associated RING-CH 1; MHCII, Major histocompatibility complex class II; MoDC, Monocyte-derived dendritic cell; PepA, Pepstatin A; shRNA, Short hairpin RNA.

2 The relevant information from our group is in the supplemental data of reference (45). Similar HLA-DR turnover was observed in additional healthy donors (C. Prevosto, A.M. Lipinska, M.J.D., R.B., unpublished data).

3 A time course of fractional synthesis in Priess cells was previously published (104). Protein fractional synthesis rates in continuously-growing cell cultures under steady-state conditions comprise two additive components: new synthesis need to (i) to maintain protein levels per cell as cells divide, and (ii) to replace proteins lost to turnover. In the Priess labelling experiments (repeated three times with similar results), concurrent measurement of cell growth rates showed that the observed fractional synthesis was entirely explained by cell growth (R.B. and A.D.R., unpublished data). We estimated that the turnover half-life must have been at least 60 hours to explain that there was no detectable difference between the fractional synthesis and cell growth rates.

4 Confirmed by N.S. and R.B. (unpublished data).

