## Supporting Information file for "Quality control of HLA-DR molecules by the lysosomal aspartyl protease, cathepsin D"

**Includes:**

**Table S1:** Mapping of peptides identified in sDR0402 CatD digestion experiments.

**Tables S2-S4** and supporting text: Prediction of CatD cleavage from primary sequence.

**Table S5:** Conservation of αF54 among human and murine MHCII α chains.

**Fig. S1:** Flow cytometry gating strategy and examples of staining.

**Fig. S2:** Variables influencing the PepA effect on DR levels.

**Fig. S3:** Example of PepA effect on DR fluorescent staining in immature MoDCs.

**Fig. S4:** PepA effect on MHCII protein levels in murine APCs.

**Fig. S5:** RT-PCR analysis of CTSD and CTSE mRNA levels in KG-1 and Priess cells.

**Fig. S6:** Peptide maps of smaller CatD fragments including β-chain peptides.

### Table S1. Mapping of peptides detected in CatD-digested sDR0402 samples[a]

**Alpha chain**

```
        1
DRA     IKEEHVIIQA EFYLNPDQSG EFMFDFDGDE IFHVDMAKKE TVWRLEEFGR
alpha   IKEEHVIIQA_EFYLNPDQSG_EFMFDFDGDE_IFHVDMAK

D1

D2                                               KE_TVWR

D3                                                      LEEFGR_

       51
DRA     FASFEAQGAL ANIAVDKANL EIMTKRSNYT PITNVPPEVT VLTNSPVELR
alpha   FASFEAQGAL_ANIAVDKANL_EIMTK

D1      FASFEAQGAL_ANIAVDKANL_EIMTK
            EAQGAL_ANIAVDK
           /\

D2      FASFEAQGAL_ANIAVDKANL_EIMTK                    TNSPVELR_
                                                     /\

D3      FASFEAQGAL_ANIAVDK                             TNSPVELR
       _F                                            /\
/\

      101
DRA     EPNVLICFID KFTPPVVNVT WLRNGKPVTT GVSETVFLPR EDHLFRKFHY
alpha   EPNVLICFID_K             NGKPVTT_GVSETVFLPR       KFHY_

D1      EPNVLICFID_K             NGKPVTT_GVSETVFLPR        FHY_

D2     _EPNVLICFID_K             NGKPVTT_GVSETVFLPR        FHY_

D3      EPNVLICFID_K             NGKPVTT_GVSETVFLPR        FHY_


      151
DRA     LPFLPSTEDV YDCRVEHWGL DEPLLKHWEF DAPSPLPETT EN
alpha  _LPFLPSTEDV_YDCRVEHWGL_DEPLLKHWEF_DAPSPLPETT_EN

D1     _LPFLPSTEDV_YDCRVEHWGL_DEPLLKHWEF_DAPSPLPETT_EN

D2     _LPFLPSTEDV_YDCRVEHWGL_DEPLLKHWEF_DAPSPLPETT_EN

D3     _LPFLPSTEDV_YDCRVEHWGL_DEPLLK_H
                       VEHWGL_DEPLLK
                       VEHWGL_DEPLL
                                  ?? ??
```

Cont'd...

Table S1 cont’d

**Beta chain**

```
          1
DRB1      GDTRPRFLEQ VKHECHFFNG TERVRFLDRY FYHQEEYVR FDSDVGEYR A
Beta            FLEQ_VK                  Y_FYHQEEYVR FDSDVGEYR A_

D1              FLEQ_VK                  Y_FYHQEEYVR FDSDVGEYR A_

D2              FLEQ_VK                  Y_FYHQEEYVR FDSDVGEYR A_
                                                               A_

D3              FLEQ_VK                  Y_FYHQEEYVR FDSDVGEYR A_
                                                               A_

         51
DRB1      VTELGRPDAE YWNSQK DILE DERAAVDTYC RHNYGVVESF TVQRRVYPEV
Beta      VTELGRPDAE_YWNSQK DILE_DERAAVDTYC_RHNYGVVESF_TVQRRVYPEV_

D1        VTELGRPDAE_YWNSQK         AAVDTYC_RHNYGVVESF_TVQRRVYPEV_

D2        VTELGRPDAE_YWNSQK         AAVDTYC_RHNYGVVESF_TVQRRVYPEV_
          VTELGRP
                ??

D3        VTELGRPDAE_YWNSQK         AAVDTYC_RHNYGVVESF_TVQRRVYPEV_
          VTELGRP
                ??

        101
DRB1      TVYPAKTQPL QHHNLLVCSV NGFYPGSIEV RWFRNGQEEK TGVVSTGLIQ
Beta      TVYPAKTQPL_QHHNLLVCSV_NGFYPGSIEV_R

D1        TVYPAK

D2        TVYPAK

D3        TVYPAK


        151
DRB1      NGDWTFQTLV MLETVPRSGE VYTCQVEHPS LTSPLTVEWR ARSESAQSKT
Beta                        SGE_VYTCQVEHPS_LTSPLTVEWR   SESAQSKT_

D1                          SGE_VYTCQVEHPS_LTSPLTVEWR
                            SGE_VYTCQVEH
                                       ??

D2                          SGE_VYTCQVEHPS_LTSPLTVEWR

D3                          SGE_VYTCQVEHPS_LTSPLTVEWR

        201
DRB1      PPPEPET
Beta     _PPPEPET                                           Cont’d…
```

[a] Table S1 key

| | |
|---|---|
| Target sequence (gaps every 10 residues) | ABC DEF |
| Tryptic fragment coverage | ABC_DEF…K/R |
| Fragment-specific non-tryptic peptide | **XYF** |
| Plausible CatD cleavage | **XYF** or **ABC**<br>/\ /\ |
| Not a plausible CatD cleavage | **XYH** or **DEF**<br>?? ?? |
| Glycosylation site | NxT |
| Beta KT3 epitope tag | **TPPPEPET** |

**Supporting text: Calculation of CatD cleavage preferences from substrate amino acid sequence and MEROPS specificity matrix.**

**Table S2** shows a specificity matrix retrieved from the MEROPS entry for CatD (access date: 29/10/2012)

**Table S2: Specificity matrix**

| Amino Acid | P4 | P3 | P2 | P1 | P1' | P2' | P3' | P4' |
|---|---|---|---|---|---|---|---|---|
| A | 52 | 30 | 64 | 32 | 69 | 92 | 51 | 57 |
| C | 12 | 7 | 21 | 8 | 11 | 12 | 11 | 5 |
| D | 41 | 56 | 58 | 31 | 35 | 37 | 52 | 51 |
| E | 47 | 72 | 80 | 29 | 43 | 67 | 69 | 62 |
| F | 64 | 26 | 19 | 163 | 93 | 15 | 17 | 17 |
| G | 39 | 42 | 20 | 17 | 7 | 44 | 47 | 69 |
| H | 6 | 4 | 8 | 0 | 0 | 2 | 3 | 7 |
| I | 39 | 61 | 50 | 0 | 87 | 38 | 42 | 25 |
| K | 29 | 28 | 31 | 5 | 29 | 63 | 65 | 71 |
| L | 105 | 80 | 51 | 333 | 92 | 50 | 94 | 44 |
| M | 17 | 24 | 20 | 27 | 38 | 12 | 22 | 26 |
| N | 20 | 18 | 40 | 4 | 20 | 17 | 36 | 33 |
| P | 42 | 24 | 3 | 1 | 3 | 2 | 11 | 52 |
| Q | 21 | 39 | 19 | 6 | 11 | 45 | 37 | 42 |
| R | 27 | 20 | 28 | 3 | 2 | 20 | 6 | 2 |
| S | 30 | 47 | 49 | 9 | 33 | 53 | 51 | 62 |
| T | 40 | 48 | 48 | 11 | 24 | 44 | 50 | 43 |
| V | 70 | 77 | 109 | 3 | 88 | 115 | 65 | 46 |
| W | 7 | 7 | 4 | 25 | 7 | 0 | 3 | 1 |
| Y | 22 | 24 | 15 | 32 | 44 | 9 | 6 | 16 |
| Total no. at position | 730 | 734 | 737 | 739 | 736 | 737 | 738 | 731 |

Dividing the number of occurrences of each amino acid by the total number of occurrences at each position, and dividing by 5% (default expectation value) yields the preference matrix in **Table S3**.Next, the amino acid sequence of interest is used to look up the cleavage preference value for each amino acid if present at each position between P4 and P4'. Lastly, the score predicting CatD cleavage after each residue (taken as P1) is computed by multiplying the look-up values for the eight positions flanking the relevant scissile bond. **Table S4** shows an example of these computations for the DR$\alpha$ sequence 49-59, centered around the empirically observed initial cleavage site C-terminal to F54. The look-up values used to compute the score for cleavage after $\alpha$F54 are highlighted in yellow. The values for the entire sequence are graphed in Fig. 3C of the main manuscript.

**Table S3: Preference matrix**

| Amino Acid | P4 | P3 | P2 | P1 | P1' | P2' | P3' | P4' |
|---|---|---|---|---|---|---|---|---|
| A | 1.42 | 0.82 | 1.74 | 0.87 | 1.88 | 2.50 | 1.38 | 1.56 |
| C | 0.33 | 0.19 | 0.57 | 0.22 | 0.30 | 0.33 | 0.30 | 0.14 |
| D | 1.12 | 1.53 | 1.57 | 0.84 | 0.95 | 1.00 | 1.41 | 1.40 |
| E | 1.29 | 1.96 | 2.17 | 0.78 | 1.17 | 1.82 | 1.87 | 1.70 |
| F | 1.75 | 0.71 | 0.52 | 4.41 | 2.53 | 0.41 | 0.46 | 0.47 |
| G | 1.07 | 1.14 | 0.54 | 0.46 | 0.19 | 1.19 | 1.27 | 1.89 |
| H | 0.16 | 0.11 | 0.22 | 0.00 | 0.00 | 0.05 | 0.08 | 0.19 |
| I | 1.07 | 1.66 | 1.36 | 0.00 | 2.36 | 1.03 | 1.14 | 0.68 |
| K | 0.79 | 0.76 | 0.84 | 0.14 | 0.79 | 1.71 | 1.76 | 1.94 |
| L | 2.88 | 2.18 | 1.38 | 9.01 | 2.50 | 1.36 | 2.55 | 1.20 |
| M | 0.47 | 0.65 | 0.54 | 0.73 | 1.03 | 0.33 | 0.60 | 0.71 |
| N | 0.55 | 0.49 | 1.09 | 0.11 | 0.54 | 0.46 | 0.98 | 0.90 |
| P | 1.15 | 0.65 | 0.08 | 0.03 | 0.08 | 0.05 | 0.30 | 1.42 |
| Q | 0.58 | 1.06 | 0.52 | 0.16 | 0.30 | 1.22 | 1.00 | 1.15 |
| R | 0.74 | 0.54 | 0.76 | 0.08 | 0.05 | 0.54 | 0.16 | 0.05 |
| S | 0.82 | 1.28 | 1.33 | 0.24 | 0.90 | 1.44 | 1.38 | 1.70 |
| T | 1.10 | 1.31 | 1.30 | 0.30 | 0.65 | 1.19 | 1.36 | 1.18 |
| V | 1.92 | 2.10 | 2.96 | 0.08 | 2.39 | 3.12 | 1.76 | 1.26 |
| W | 0.19 | 0.19 | 0.11 | 0.68 | 0.19 | 0.00 | 0.08 | 0.03 |
| Y | 0.60 | 0.65 | 0.41 | 0.87 | 1.20 | 0.24 | 0.16 | 0.44 |

**Table S4: Example of cleavage susceptibility score calculation**

| DR alpha | | | | | | | | | | | |
|---|---|---|---|---|---|---|---|---|---|---|---|
| **AA no.** | **49** | **50** | **51** | **52** | **53** | **54** | **55** | **56** | **57** | **58** | **59** |
| **AA** | **G** | **R** | **F** | **A** | **S** | **F** | **E** | **A** | **Q** | **G** | **A** |
| **Preference matrix lookup values** | | | | | | | | | | | |
| **P4** | 1.07 | 0.74 | 1.75 | 1.42 | 0.82 | 1.75 | 1.29 | 1.42 | 0.58 | 1.07 | 1.42 |
| **P3** | 1.14 | 0.54 | 0.71 | 0.82 | 1.28 | 0.71 | 1.96 | 0.82 | 1.06 | 1.14 | 0.82 |
| **P2** | 0.54 | 0.76 | 0.52 | 1.74 | 1.33 | 0.52 | 2.17 | 1.74 | 0.52 | 0.54 | 1.74 |
| **P1** | 0.46 | 0.08 | 4.41 | 0.87 | 0.24 | 4.41 | 0.78 | 0.87 | 0.16 | 0.46 | 0.87 |
| **P1'** | 0.19 | 0.05 | 2.53 | 1.88 | 0.90 | 2.53 | 1.17 | 1.88 | 0.30 | 0.19 | 1.88 |
| **P2'** | 1.19 | 0.54 | 0.41 | 2.50 | 1.44 | 0.41 | 1.82 | 2.50 | 1.22 | 1.19 | 2.50 |
| **P3'** | 1.27 | 0.16 | 0.46 | 1.38 | 1.38 | 0.46 | 1.87 | 1.38 | 1.00 | 1.27 | 1.38 |
| **P4'** | 1.89 | 0.05 | 0.47 | 1.56 | 1.70 | 0.47 | 1.70 | 1.56 | 1.15 | 1.89 | 1.56 |
| **Multiplication of lookup values yields a score for cleavage potential after each residue:** | | | | | | | | | | | |
| **Score** | 0.03 | 0.16 | 14.18 | 0.28 | 1.62 | 46.43 | 3.36 | 0.65 | 1.83 | 0.79 | 2.96 |

The method can be applied to any protease with a substrate specificity matrix in the MEROPS database, updated with more recent versions of the specificity matrices, or refined by using a more sophisticated way of computing the preference matrix. The approach does not take into account secondary or higher-order protein structure and the resultant constraints on the accessibility of the cleavage site.

**Table S5. Sequence alignment near the extended peptide-flanking α-chain strand of HLA class II molecules[a]**

<table>
<tr><th>Residue</th><th>41</th><th>51</th><th></th><th>61 70</th><th>Score[c]</th></tr>
<tr><th>Sec struct</th><th>BBBTT33333</th><th>3SSS</th><th>SSHHHH</th><th>HHHHHHHHHH</th><th></th></tr>
<tr><td>DRA</td><td>TVWRLEEFGR</td><td>FASF</td><td>|EAQGAL</td><td>ANIAVDKANL</td><td>46.4</td></tr>
<tr><td>DPA1*01:03</td><td>TVWHLEEFGQ</td><td>AFSF</td><td>|EAQGGL</td><td>ANIAILNNNL</td><td>32.7</td></tr>
<tr><td>DPA1*02:01</td><td>---------R</td><td>----</td><td>|------</td><td>----------</td><td>32.7</td></tr>
<tr><td>DQA1*01:01[b]</td><td>TAWRWPEFSK</td><td>FGGF</td><td>|DPQGAL</td><td>RNMAVAKHNL</td><td>0.47</td></tr>
<tr><td>DQA1*02:01</td><td>-V-KL-L-HR</td><td>LR.-</td><td>|---F--</td><td>T-I--L----</td><td>0.13</td></tr>
<tr><td>DQA1*03:01</td><td>-V-QL-L-RR</td><td>-RR-</td><td>|---F--</td><td>T-I--L----</td><td>0.08</td></tr>
<tr><td>DQA1*04:01</td><td>-V-CL-VLRQ</td><td>-R.-</td><td>|---F--</td><td>T-I--T----</td><td>0.03</td></tr>
<tr><td>H2-Eα[d]</td><td>TIWRLEEFAK</td><td>FASF</td><td>|EAQGAL</td><td>ANIAVDKANL</td><td>46.4</td></tr>
<tr><td>H2-Aα[d/g7]</td><td>TVWRLPEFGQ</td><td>LILF</td><td>|EPQGGL</td><td>QNIAAEKHNL</td><td>3.50</td></tr>
<tr><td>H2-Aα[b]</td><td>---M------</td><td>-AS-</td><td>|D-----</td><td>----VV----</td><td>1.35</td></tr>
<tr><td>H2-Aα[k]</td><td>---M----A-</td><td>-RR-</td><td>|------</td><td>----TG----</td><td>0.63</td></tr>
<tr><td>H2-Aα[u]</td><td>-I-M----A-</td><td>-RS-</td><td>|D-----</td><td>----TG----</td><td>0.90</td></tr>
<tr><td>H2-Aα[s]</td><td>-I-M------</td><td>-TS-</td><td>|D-----</td><td>----TG-YT-</td><td>2.16</td></tr>
<tr><td>H2-Aα[r]</td><td>---M------</td><td>-TS-</td><td>|D-----</td><td>----VV----</td><td>2.16</td></tr>
<tr><td>H2-Aα[q]</td><td>---M------</td><td>-TS-</td><td>|D-----</td><td>----TG----</td><td>2.16</td></tr>
<tr><td>H2-Aα[f]</td><td>----------</td><td>-TS-</td><td>|D-----</td><td>-E--TG----</td><td>2.16</td></tr>
</table>

Notes:

[a] For each locus, the reference sequence is shown in full single-letter code, and relevant allelic variants with dashes for identity and dots for deletions. Putative or (for DR) actual cleavage at "|". CatD cleavage scores computed as in Materials and Methods. Secondary structure (from RCSB PDB 5NI9 Protein Feature View): B = beta sheet, T = turn, 3 = $3_{10}$ helix, S = strand (hydrogen bonding with peptide), A = alpha helix.

[b] DQA1 and H2-A alleles were selected to reflect a range of polymorphisms in the vicinity of the cleavage site.

[c] Score = Predicted CatD susceptibility score at αF54. In some alleles, other nearby residues show local maxima in CatD susceptibility scores: Aα[d/g7] scores 24 at L53. More distant, outside the strand, Aα[f] scores 14 at E62; the other Aα alleles score 5.8-17 at F48; DQA1*01:01 scores 20 at M63; the other DQA1 alleles score 5-6 at N62.

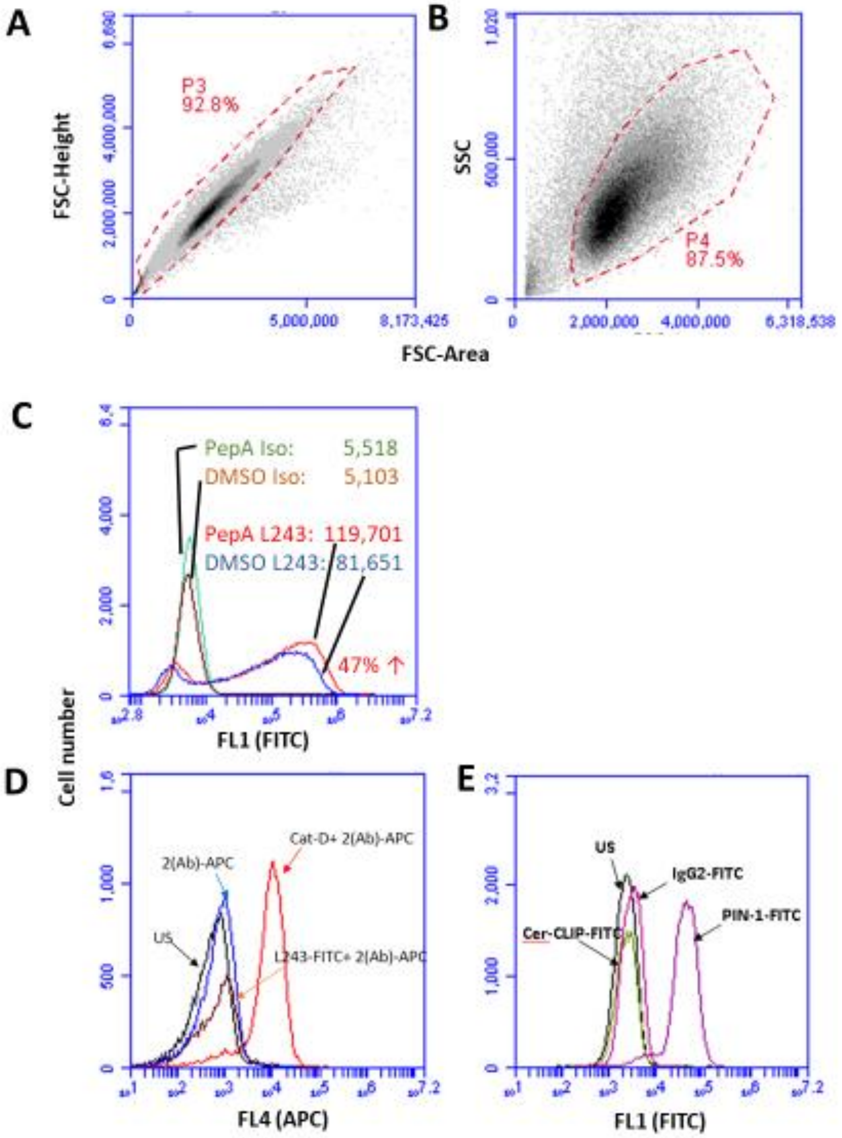


**Fig. S1. Representative flow cytometry plots of fixed, permeabilized KG-1 cells.**

All analyses were performed on Accuri C6 flow cytometer.

A and B, Gating strategy for doublet exclusion by plotting FSC-height vs. area (A) and FSC/SSC gate for identification of intact cells (B).

C, Comparison of isotype control and anti-DR (L243) staining between vehicle- (DMSO) or PepA-treated, fixed and permeabilized KG-1 cells. Heterogeneous anti-DR staining was observed despite thorough mixing of staining reactions, as observed by others (109).

In our hands, the staining profiles remained stable on time scales of months, but the degree of heterogeneity varied between independent subcultures over several years of study.

D, Specific staining with rabbit anti-CatD antibody and demonstration of lack of crossreactivity with mouse IgG (L243).

E, Staining for Ii (PIN.1) and CLIP (CerCLIP.1) vs. unstained and isotype controls.

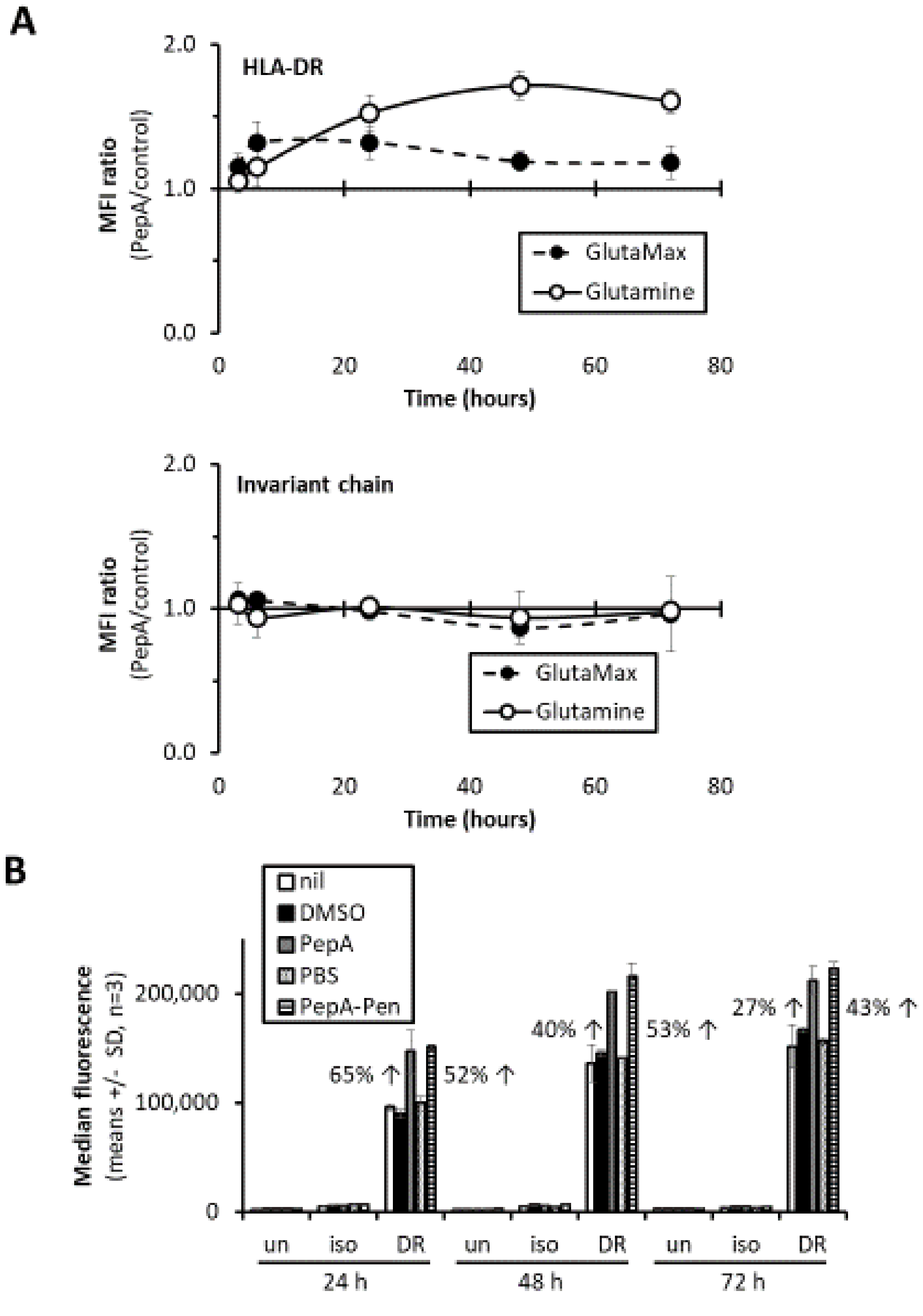


**Fig. S2. Variables affecting PepA rescue of HLA-DR molecules in KG-1 cells.**

A, Earlier, transient peak PepA effect on HLA-DR levels in KG-1 cells grown in GlutaMax media. KG-1 cells were grown normally (glutamine) or switched to GlutaMax media for at least 5 doublings, then cultured with 20 μM PepA or vehicle control (DMSO), fixed and permeabilized, and stained for HLA-DR (L243-FITC, *top*) or Ii (PIN.1-FITC), *bottom*. Shown are FL1 median fluorescence ratios (mean ± SD, n = 3) for staining of PepA-treated vs. control cells. The SDs for the ratios were calculated using the error propagation formula, $CV_{x/y} = \sqrt{(CV_x^2 + CV_y^2)}$, which is adequate for CVs ≤ 10% generally obtained here (110). For anti-DR MFIs in GlutaMax media, the PepA effect was statistically significant (1 df, $F = 71.4$, $p < 0.001$); the Ii effect was not (1 df, $F = 0.124$, $p = 0.729$; 2-way ANOVA with time as a second fixed factor).

B, Rescue of DR molecules by PepA-penetratin. KG-1 cells were grown for up to 3 days in the presence of no additives, 20 mM PepA, DMSO (vehicle for PepA), 10 μM PepA-penetratin (PepA-pen) or PBS (vehicle for PepA-pen), with flow cytometric analysis for total (surface plus intracellular) DR as in Fig. 1B. By 2-way ANOVA (with time as the second fixed factor and Tukey's HSD post-test), the effect of PepA-penetratin was statistically significant (2 df, $F = 134$, $p < 0.001$ for the overall treatment effect) when compared to untreated or vehicle controls ($p < 0.001$ for each comparison).

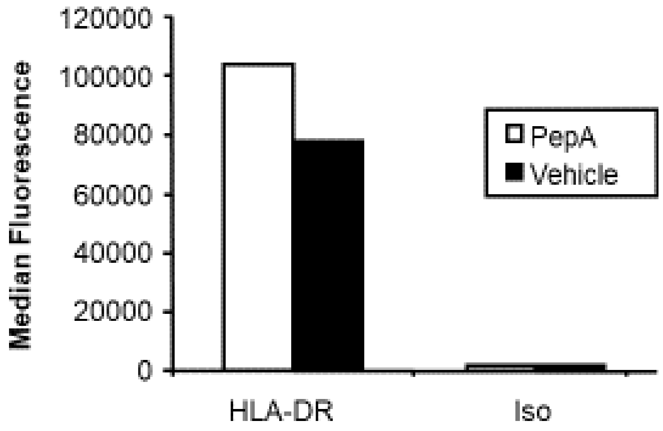


**Fig. S3. Representative example of PepA effect on HLA-DR expression in immature MoDCs.** Median fluorescence intensities (individual measurements) are shown for representative immature MoDCs from healthy donors, stained for HLA-DR (L243-FITC) and isotype controls (iso).

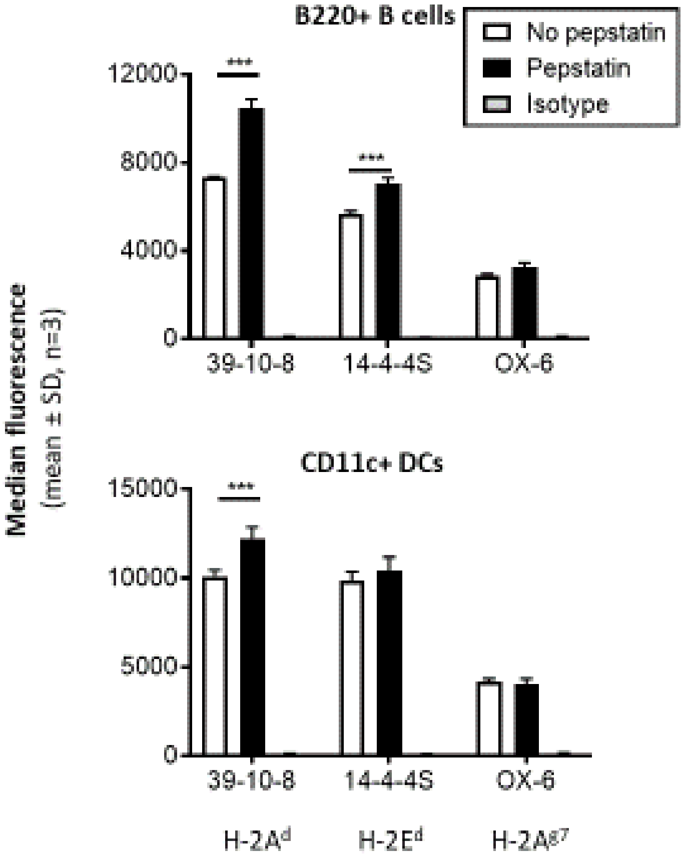


**Fig. S4. Effect of PepA on the expression of murine MHC class II proteins in splenic APCs.** Splenocytes from (Balb/c × NOD) F1 mice were treated with PepA for 24 hours, then stained for lineage markers of B cells (B220) and DCs (CD11c) and with the indicated anti-H-2A$^{d}$, E$^{d}$ or A$^{g7}$ mAbs. MHCII staining was determined after gating for each cell lineage. In B cells, 2-way ANOVA of MHCII staining data showed significant differences in staining between the different MHCII antibodies, a significant effect of PepA treatment, and a significant interaction between the two factors ($p < 0.0001$ each). In DCs, all three effects were also significant ($p = 0.006$ or less). In both graphs, *** indicates a significant PepA effect by Sidak post-test ($p < 0.0006$ or less).

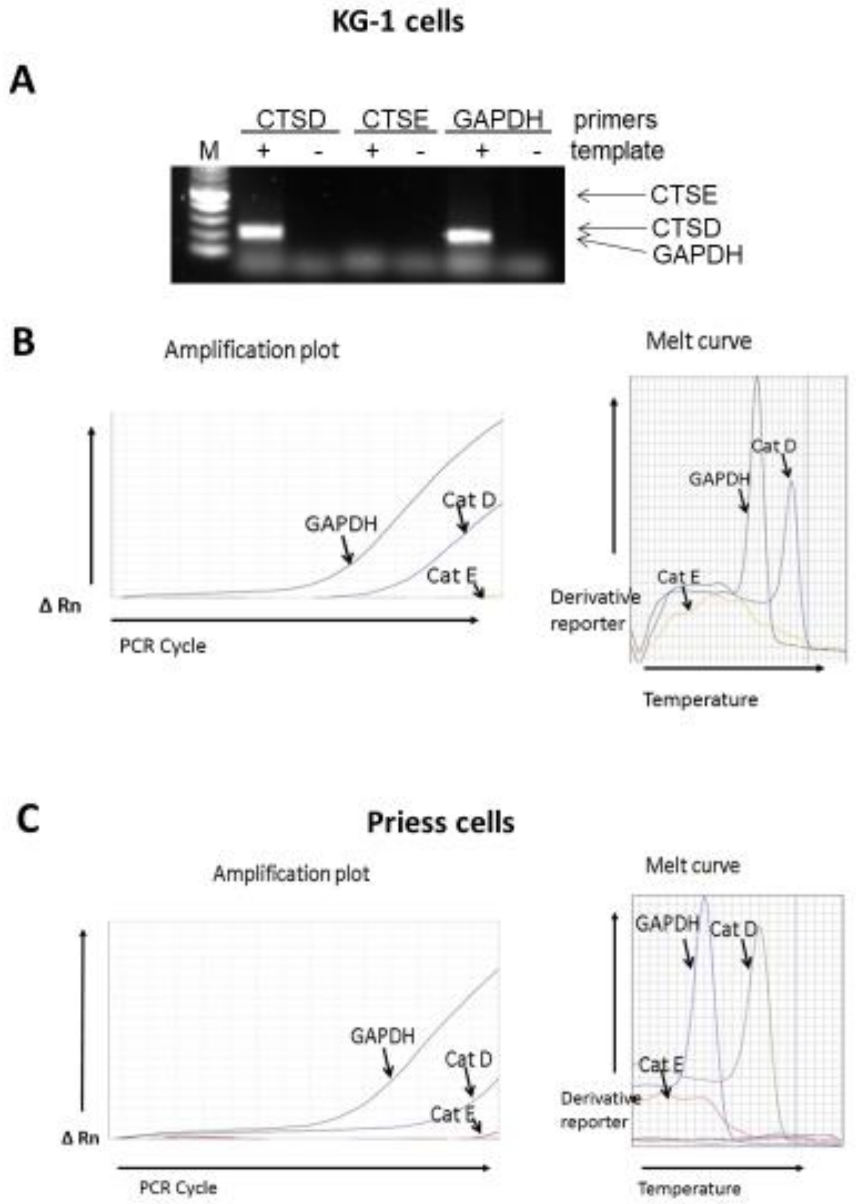


**Fig. S5. RT-PCR analysis of CTSD and CTSE transcripts, coding for CatD and CatE respectively.**

A, agarose gel electrophoresis of PCR products amplified from KG-1 cDNA. Primers for CTSD, CTSE and GAPDH were used. No-template controls are shown. Molecular weight marker (100 bp ladder, lane M) and the expected sizes of amplicons are indicated.

B and C, real-time quantitative RT-PCR analysis for the indicated mRNAs in KG-1 (B) and Priess (C) cells, using the SYBR Green method. Amplification plots (*left* panels, Δ Rn = SYBR Green fluorescence) and melting curves (*right* panels, y axis = d(fluorescence/dT)) are shown. Similar results were obtained in three independent experiments.

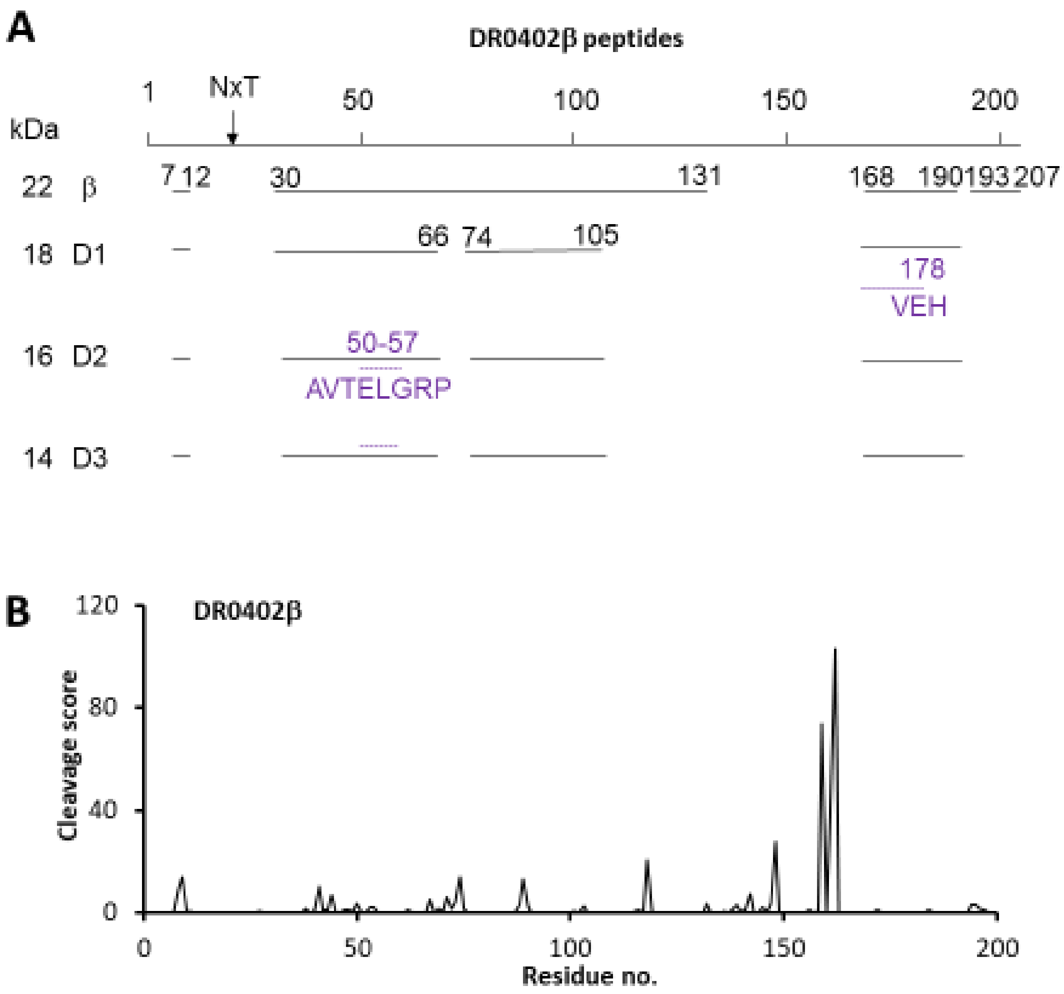


**Fig. S6. Mapping of peptides in the D1-D3 fragments of Fig. 3A to the amino acid sequence of soluble recombinant DR0401 β chain.** See legend to Fig. 3, B and C for details and Table S1 for sequences.
